# Cellular Chemical Dynamics Governing Signal Transduction and Adaptive Gene Expression: Beyond Classical Kinetics

**DOI:** 10.64898/2026.02.13.705865

**Authors:** Jinhyung Kim, Seohyun Kim, Sungho Jang, Seong Jun Park, Sanggeun Song, Kumyoung Jeung, Gyoo Yeol Jung, Ji-Hyun Kim, Hye Ran Koh, Jaeyoung Sung

**Affiliations:** Global Science Research Center for Systems Chemistry, Chung-Ang University, Seoul 06974, Korea; Creative Research Initiative Center for Chemical Dynamics in Living Cells, Chung-Ang University, Seoul 06974, Korea; Department of Chemistry, Chung-Ang University, Seoul 06974, Korea; Department of Bioengineering and Nano-Bioengineering, Incheon National University, 119 Academy-ro, Yeonsu-gu, Incheon 22012, Korea; Division of Bioengineering, College of Life Sciences and Bioengineering, Incheon National University, 119 Academy-ro, Yeonsu-gu, Incheon 22012, Korea; Research Center for Bio Materials and Process Development, Incheon National University, 119 Academy-ro, Yeonsu-gu, Incheon 22012, Korea; Department of Physics and Astronomy and Center for Theoretical Physics, Seoul National University, Seoul 08826, Korea; Department of Chemical and Biomolecular Engineering, College of Chemistry, University of California, Berkeley, Berkeley, CA 94720, USA; Chemical Science Division, Lawrence Berkeley National Laboratory, Berkeley, CA 94720, USA; Division of Interdisciplinary Bioscience and Bioengineering, Pohang University of Science and Technology, 77 Cheongam-ro, Nam-gu, Pohang, Gyeongbuk, 37673, Korea; Department of Chemical Engineering, Pohang University of Science and Technology, 77 Cheongam-ro, Nam-gu, Pohang, Gyeongbuk, 37673, Korea

## Abstract

Cellular adaptation is inherently nonstationary processes with complex stochastic dynamics^1-5^. Despite remarkable progress in quantitative biology^6-11^, a quantitative understanding of the cell adaptation dynamics in terms of the underlying cellular network remains elusive. Here, we present the next-generation chemical dynamics model and theory for cellular networks, providing an effective, quantitative description of the adaptive gene expression dynamics in living cells responding to external stimuli. Unlike conventional kinetics, chemical dynamics of cellular network modules are characterized by their reaction-time distributions, rather than by rate coefficients^12^. For a general model of cell signal transduction and adaptive gene expression, we derive exact analytical expressions for the time-dependent mean and variance of protein numbers produced in response to external stimuli, validated by accurate stochastic simulations. These results provide a unified, quantitative explanation of the stochastic responses of diverse *E. coli* genes to antibiotic stress and transcriptional induction. Our analysis reveals existence of a general quadratic relationship between the mean and variance of activation times across diverse genes. The gene activation process influences transient dynamics of downstream protein levels, but not their steady-state levels. In contrast, post-translational maturation process affects both transient dynamics and steady-state variability of mature protein levels. This finding indicates that the gene expression variability measured by fluorescent reporter proteins depends on the maturation time of the reporters. This work suggests a new direction for the development of digital twins of living cells.

---

Adaptation to environmental change is a defining property of life^13-16^. It occurs through a complex, hierarchical network of intracellular chemical processes, which can be categorized into several functional modules. The chemical dynamics of these modules govern the distribution of time-lag between the onset of an environmental perturbation and the ensuing adaptive cell response ^17-20^. Through evolution, life forms have been optimizing the chemical dynamics of their regulatory reaction networks to ensure timely adaptation and survival under environmental changes^21,22^. An important goal of modern quantitative biology is to achieve a quantitative understanding of complex dynamics of these regulatory reaction networks underlying adaptation. Modern fluorescence imaging techniques have enabled real-time monitoring of adaptation dynamics of individual cells under various external stimuli^23-26^. Pioneering studies investigated the adaptation dynamics of *S. cerevisiae* under periodic osmotic stress^27^ and the damped oscillatory gene-expression dynamics of *p53* and NF-κB in mammalian cells responding to DNA damage and inflammatory signals^28-32^. More recent work has collectively tracked the expression dynamics of diverse *E. coli* genes for hundreds of individual cells subjected to antimicrobial stress^4,16^. These experimental studies demonstrated that adaptive gene expression is a gene-specific, nonstationary process, whose dynamics exhibits significant cell-to-cell variation. Despite remarkable progress in network biology and quantitative biology^33,34^, achieving a quantitative understanding of stochastic cell-adaptation dynamics remains a formidable challenge. This difficulty arises because the reaction-network modules underlying cellular adaptations have far more complex dynamics than assumed in the conventional theories based on chemical kinetics or chemical master equations^35,36^.

In this work, we address this issue by presenting a new type of chemical-dynamics model for signal transduction and adaptive gene expression. The underlying reaction network is decomposed into key functional modules, including signal sensing and propagation leading to gene activation, gene expression, protein maturation, and protein degradation. Chemical dynamics of each module is characterized by its reaction-time distribution (RTD), or the distribution of the time required to complete a single modular process. The use of an RTD enables a more accurate characterization of the chemical dynamics of an intracellular reaction-network module^12^, which may comprise numerous *vibrant* enzymatic reactions, than conventional models that merely assign a single rate coefficient to each module^37^.

For our model, we derive exact analytical expressions for the time-dependent mean and variance of the number of proteins produced in response to gene-activating external stimuli. The correctness of these results is verified through accurate stochastic simulations. Our results provide a unified, quantitative explanation for the time-dependent mean and variance of 23 protein levels in *E. coli*, expressed in response to two stress signals and one gene-induction signal. From these analyses, we extracted the RTDs of gene-activation processes for various chromosomal *E. coli* genes and uncovered a quadratic relationship between the mean and variance of gene-activation times across diverse chromosomal genes activated by antimicrobial stresses.

Our analyses reveal that signal-transduction processes preceding target gene expression modulate the transient dynamics of the mean and variance of mature protein levels but leave their steady-state values unchanged. In contrast, protein-maturation processes following the gene expression influence both the temporal behavior and the steady-state variance of mature protein levels. Quantitative predictions are made for how gene expression dynamics, variability, and correlations measured using fluorescent reporter proteins depend on the maturation-dynamics of the reporters, an effect previously unrecognized. The present theory also enables the disentanglement of the intrinsic temporal profiles and steady-state variability of adaptive gene expression from the confounding influence of reporter-maturation dynamics. Together with modern high-throughput measurements of cellular dynamics and advances in machine-learning techniques, this work suggests a promising new direction for quantitative understanding of cell dynamics and the systematic development of digital twins of living cells.

## Cell Adaptation Model

We consider a clonal population of cells that express a set of genes to adapt to stepwise environmental changes. The underlying reaction networks can be decomposed into signal-sensing and propagation leading to gene activation, gene-expression, protein-maturation, and degradation modules (Fig 1a). Each module can be further divided into smaller modules when relevant information is available. For example, the gene-expression module can be decomposed into the first gene-expression-burst submodule and the subsequent burst submodules, because the first burst exhibits distinct dynamics from the rest.

**Figure 1.**
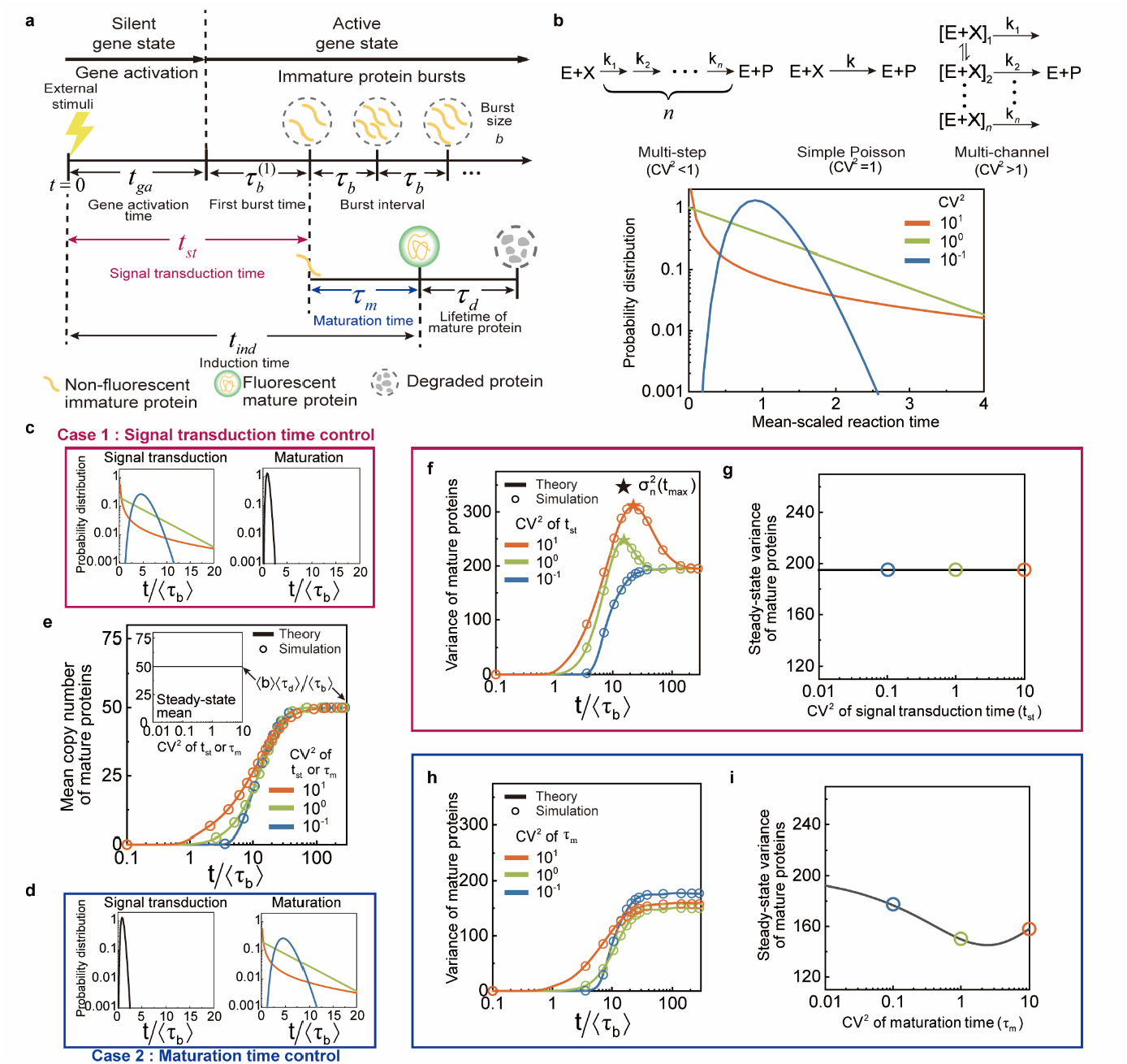
Signal-induced adaptive gene expression model and distinct roles of signal transduction time and maturation time in mature-protein dynamics and steady-state statistics. **a**, Schematic of the signal-induced adaptive gene expression model (see Supplementary Note 3 for details). **b**, Relationship between the shape of the reaction-time distribution and the underlying kinetic scheme: a simple Poisson process (*CV*^2^ = 1), a multi-step sub-Poisson process (*CV*^2^ < 1), and a multi-channel super-Poisson process (*CV*^2^ > 1). **c–i**, Effects of signal transduction time (*t*_*st*_) and protein maturation time (*τ*_*m*_) on mature-protein dynamics and steady-state statistics. (**c**) Case 1: varying the *CV*^2^ of *t*_*st*_ with ⟨*t*_*st*_ ⟩ ⟨*τ*_*b*_ ⟩ = 5 and ⟨*τ*_*m*_ ⟩ ⟨*τ*_*b*_ ⟩ = 1 . (**d**) Case 2: varying the *CV*^2^ of *τ*_*m*_ with ⟨*t*_*st*_ ⟩ ⟨*τ*_*b*_ ⟩ = 1 and ⟨*τ*_*m*_ ⟩ ⟨*τ*_*b*_ ⟩ = 5 . (**e**) Mean mature-protein level, identical in both cases. The inset shows that the steady-state mean is independent of the *CV*^2^ of *t*_*st*_ or *τ*_*m*_ . (**f**) Mature protein variance and (**g**) its steady-state value in Case 1. In **f**, the variance, 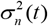, exhibits a crossover from monotonic to non-monotonic time dependence, developing a peak at *t* = *t*_max_, as the *CV*^2^ of *t*_*st*_ increases, while the steady-state variance in **g** is independent of the *CV*^2^ of *t*_*st*_ . (**h**) Mature protein variance and (**i**) its steady-state value in Case 2. In **h**, the variance increases monotonically with time, irrespective of the *CV*^2^ of *τ*_*m*_, while the steady-state variance in **i** depends non-monotonically on the *CV*^2^ of *τ*_*m*_ . In **c–i**, both *t*_*st*_ and *τ*_*m*_ are assumed to follow a gamma distribution, 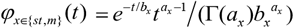. The time interval, *τ*_*b*_, between successive immature-protein burst events and the mature-protein lifetime, *τ*_*d*_, are assumed to follow an exponential distribution, 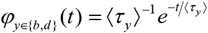 with ⟨*τ*_*d*_ ⟩ ⟨*τ*_*b*_ ⟩ = 10 . Values of the first and second moments of the burst size, *b*, are respectively given by ⟨*b*⟩ = 5 and ⟨*b*^2^ ⟩ = 35 . In **e** and **f–i**, symbols and lines represent stochastic simulation results and theoretical predictions [equations (1a) and (1b)] for the mature-protein mean and variance, respectively. Further details of the stochastic simulation method are provided in Supplementary Note 8.

The chemical dynamics of each modular network can be modeled more accurately by employing its reaction time distribution (RTD) than by using a rate coefficient. When detailed information about the mechanisms and dynamics of the reactions and molecular transport processes constituting a network module is available, the functional form of the RTD of the module can be derived^38^ (Fig. 1b for simple examples). In the absence of such information, the RTD can still be extracted from experimental data, as demonstrated below. We find that the RTD of a multistep process approaches a sub-Poisson gamma distribution as the number of steps increases (see Supplementary Note 1), whereas the RTD of a network with a single rate-determining step with cell-state-dependent fluctuations in its rate coefficient can be approximated by a super-Poisson gamma distribution (see Supplementary Note 2). Therefore, these gamma distributions, with only two parameters, can effectively characterize the RTDs of large reaction network modules, greatly reducing the number of adjustable parameters.

In our model, the signal transduction designates the composite process consisting of gene activation and the first gene expression burst processes. Accordingly, the signal transduction time is defined as the sum of the gene activation time and the first gene expression burst time. In addition, the induction time is defined as the sum of the signal transduction time and the protein maturation time. The RTDs of gene activation, the first gene expression burst, and subsequent bursts are modeled by distinct gamma distributions. In case where the subsequent bursts occur on a time scale far shorter than gene activation and the first burst, the detailed shape of the RTD of each individual burst has little effect on overall cell-adaptation dynamics and the RTD of each burst can be approximated by the simple exponential function. The maturation time distributions of fluorescent reporter proteins can be extracted from existing experimental results. More details about our model, along with our nomenclature, are presented in Fig. 1a and Supplementary Note 3.

### Analytic Results

Our model yields exact analytical expressions for the time-dependent mean, ⟨*n*(*t*)⟩, and variance, 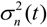, of the mature protein number (see Methods):

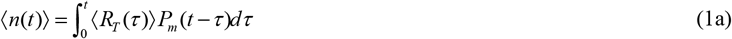

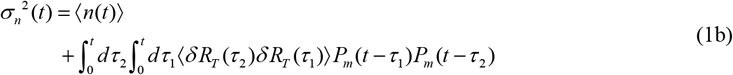

where ⟨*R*_*T*_ (*τ*)⟩ denotes the average production rate of immature proteins at time *τ*, measured from the onset of environmental change. Angular brackets denote averages taken over the trajectories of a cell population. *P*_*m*_ (*t*) denotes the probability that an immature protein generated at time 0 has completed its maturation process and survived as a mature protein at time *t*, given by

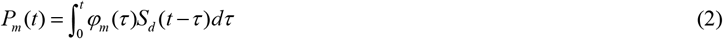

where *φ*_*m*_ (*t*) and *S*_*d*_ (*t*) designates the maturation time distribution and the survival probability of mature proteins, respectively.

As shown in equation (1b), the variance of mature protein number depends on the time-correlation function (TCF), ⟨*δ R*_*T*_ (*τ*_2_)*δ R*_*T*_ (*τ*_1_)⟩, of the fluctuations, *δ R*_*T*_ (*t*) = *R*_*T*_ (*t*) − ⟨*R*_*T*_ (*t*)⟩, in the creation rate of immature proteins. The explicit expressions of ⟨*R*_*T*_ (*t*)⟩ and ⟨*δ R*_*T*_ (*τ*_2_)*δ R*_*T*_ (*τ*_1_)⟩ are obtained in terms of signal transduction time distribution *φ*_*st*_ (*t*), protein burst interval distribution *φ*_*b*_ (*t*), and the first two moments of the burst size, i.e., the number of nascent proteins created per gene-expression burst (see Methods and Supplementary Note 4). We confirm the correctness of our analytic results using accurate stochastic simulations (Fig. 1e, f, h; Fig. 2h, i).

**Figure 2.**
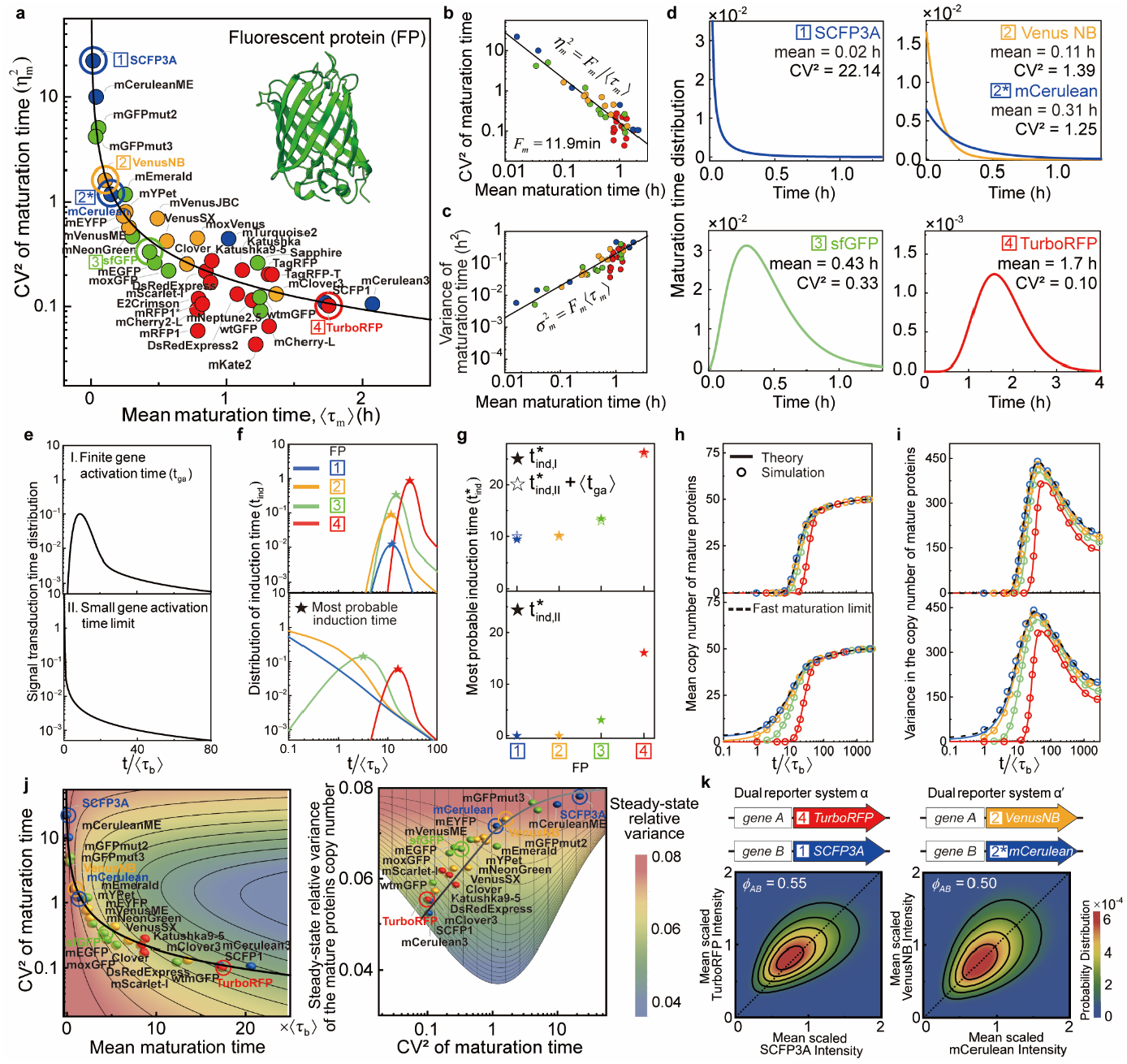
Protein maturation time distributions and their impact on mature-protein dynamics and steady-state statistics. **a**, Relation between the mean (⟨*τ* ⟩) and 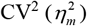 of the protein maturation time distribution extracted for fluorescent proteins emitting in the cyan, green, yellow, and red spectral ranges from translation-stopped experiments at 37 °C^43^ (Supplementary Fig. 3 and Supplementary Note 9 for details). The solid line represents a best fit of 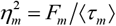 with the value of the Fano factor, *F*, given by 11.9 min. **b**, Double-logarithmic plot of the solid line in **a. c**, Linear mean-variance relationship for the protein maturation time 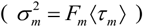. **d**, Representative maturation time distributions. **e–i**, Dependence of mature-protein dynamics on the maturation time distribution. Case I: finite gene activation time (upper); Case II: small gene activation time limit (lower). (**e**) Signal transduction time distribution. In the upper panel, signal-induced gene activation and the ensuing first protein-burst event are treated as a multi-step process and a multi-channel process, respectively, the latter reflecting heterogeneous kinetics routes leading to burst initiation; the corresponding distributions are given by gamma distributions with mean and CV^2^ given by 10⟨*τ*_*b*_ ⟩ and 0.1, and by 50⟨*τ*_*b*_ ⟩ and 21. In the lower panel, gene activation times are set to zero, while the first burst time distribution is identical to that in the upper panel. (**f**) Induction time distribution. In the upper panel, the induction time distribution is given by the convolution of the signal transduction time distribution in **e** and the maturation time distribution in **d.** (**g**) Most probable induction time. In Case I, the induction time approximately equals the sum of the induction time in Case II and the mean gene activation time (open stars). (**h**) Mean and (**i**) variance of mature protein levels. **j**, Steady-state relative variance, 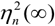, of mature protein levels as a function of the mean and CV^2^ of the maturation time distribution: the top view (left) and the right-side view (right). These plots illustrate how experimentally extracted maturation parameters map onto the theoretical landscape for 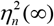. The solid lines represent the trace of the theoretical surface evaluated along the curve, 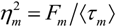, shown in **a** [see equation (N5-10) in Supplementary Note 5]. **k**, Reporter dependence of the joint distribution of observed expression levels in dual-reporter systems: SCFP3A and TurboRFP (left), and CFP and YFP (right). The correlation coefficient *ϕ*_*AB*_ between the observed expression levels becomes reporter dependent because fluorescent protein maturation selectively affects their absolute variances without altering their means and covariance, the latter reflecting the intrinsic upstream coupling between genes *A* and *B*. Throughout this figure, the distributions of the burst interval and the mature-protein lifetime, and the values of ⟨*b*⟩ and ⟨*b*^2^ ⟩ are the same as those in Fig. 1, except in **k**, where ⟨*b*^2^ ⟩ = 100 .

Our results show that the signal transduction and protein maturation processes have strong influence on the transient dynamics of the mean mature protein number but do not influence its steady-state value. The *steady-state mean* of the mature protein number is determined by the gene-expression and protein-degradation dynamics, specifically by the mean burst size ⟨*b* ⟩, the mean burst time interval ⟨*τ*_*b*_ ⟩ and the mean lifetime ⟨*τ*_*d*_ ⟩ of mature proteins, i.e., 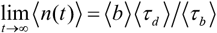 (see Methods and Supplementary Note 5). In contrast, the *temporal profile* of ⟨*n*(*t*)⟩ primarily depends on the convolution of the signal transduction time distribution, *φ*_*st*_ (*t*), and protein maturation time distribution, *φ*_*m*_ (*t*), given that individual gene expression bursts occurs on a time scale far shorter than the signal transduction and protein maturation (see Methods). Owing to the commutative property of convolution, signal transduction and protein maturation equally contribute to the time evolution of ⟨*n*(*t*) ⟩ (Fig. 1c– e).

Unlike the mean, the variance of the mature-protein number differently depends on signal transduction and protein maturation dynamics (Fig. 1f, h). The *steady-state* variance ⟨*δn*^2^ (∞)⟩ of the mature protein number is sensitive to the maturation dynamics but is independent of the signal transduction dynamics (Fig. 1g, i). ⟨ *δn*^2^ (∞) ⟩ decreases with the mean maturation time but has a nonlinear dependence on the randomness of the protein maturation time, reaching a minimum at a particular value of the maturation time randomness when the mean maturation time is fixed (Fig. 1i). In the fast maturation limit, the steady-state protein number fluctuation is completely determined by the gene expression dynamics and the protein lifetime distribution ^12,37^. On the other hand, the *transient dynamics* of the variance of the mature protein number are sensitive to signal transduction as well as protein maturation dynamics (Fig. 1f, h). The variance monotonically increases with time up to its long-time steady-state value if the signal transduction is a strongly sub-Poisson process. Otherwise, the variance can exhibit a non-monotonic time dependence, attaining a maximum before relaxing to its steady-state value, especially when the signal transduction occurs on a time scale far longer than that of the individual gene expression bursts (see Supplementary Fig. 1).

Our theory predicts that steady-state statistics, not to mention transient dynamics, of gene expression levels measured using a fluorescent reporter protein depend on the maturation time distribution of the fluorescent reporter itself. This is demonstrated for two distinct models of signal transduction in Fig. 2e. The maturation time of fluorescent reporter proteins exhibits a substantial reporter-to-reporter variation (Fig. 2a); the mean maturation time ranges from 12 minutes to 2.1 hours, and the relative variance of the maturation time ranges from 0.04 to 20. We find that the relative variance, 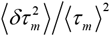, of the maturation time tends to be inversely proportional to the mean maturation time, ⟨ *τ*_*m*_ ⟩, and is approximately given by 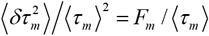 with *F*_*m*_ ≅ 11.9 min for diverse fluorescent reporter proteins (Fig. 2a, b and Supplementary Note 6). Most fluorescent reporters exhibit a unimodal maturation time distribution with the mean greater than 11.9 min and the relative variance smaller than unity. The shape of the maturation time distribution of fluorescence reporters influences the dynamics of the mature-protein copy number as well as the distribution of induction time (Fig. 2d-i). Our theory predicts that the steady-state gene expression variability, measured using a fluorescent reporter protein, increases (decreases) with the relative variance (mean) of the maturation time of the fluorescent reporter (Fig. 2j). Consequently, intrinsic noise and extrinsic noise, or correlation between gene expression levels, measured by a dual reporter system^6^ depend on the maturation time distributions of the two fluorescent reporter proteins employed in the system. Equations (1a) and (1b) enable us to quantify and disentangle the confounding effects of reporter maturation dynamics on the gene expression statistics measured using fluorescence reporters (Fig. 2k).

### Quantitative Analyses of Experimental Data

We compare our theory with three sets of experimental data on adaptive gene expression in *E. coli*, obtained under two antibiotic stress signals^4^ and one transcriptional induction signal (see Methods). The antibiotic stress-response data were obtained by employing yellow or cyan fluorescent proteins (YFP^39^/CFP^40^) expressed under various chromosomal *E. coli* promoters activated by tetracycline (TET) or trimethoprim (TMP) stress signals in Ref. ^4^ (Fig. 3a). On the other hand, the transcriptional induction-response data were obtained by employing super-folder GFP (sfGFP) expressed from three *E. coli* plasmids with controllable transcriptional efficiencies, which can be induced by isopropyl β-D-1-thiogalactopyranoside (IPTG) (see Methods). Translation efficiencies of these plasmids were modulated by adjusting the sequences of their 5′-untranslated regions (5′-UTRs)^41,42^ (Fig. 3d). For these systems, we obtained experimental datasets for the time-dependent mean and variance of the mature fluorescent protein reporters. These data could not be quantitatively explained by previous models.

**Figure 3.**
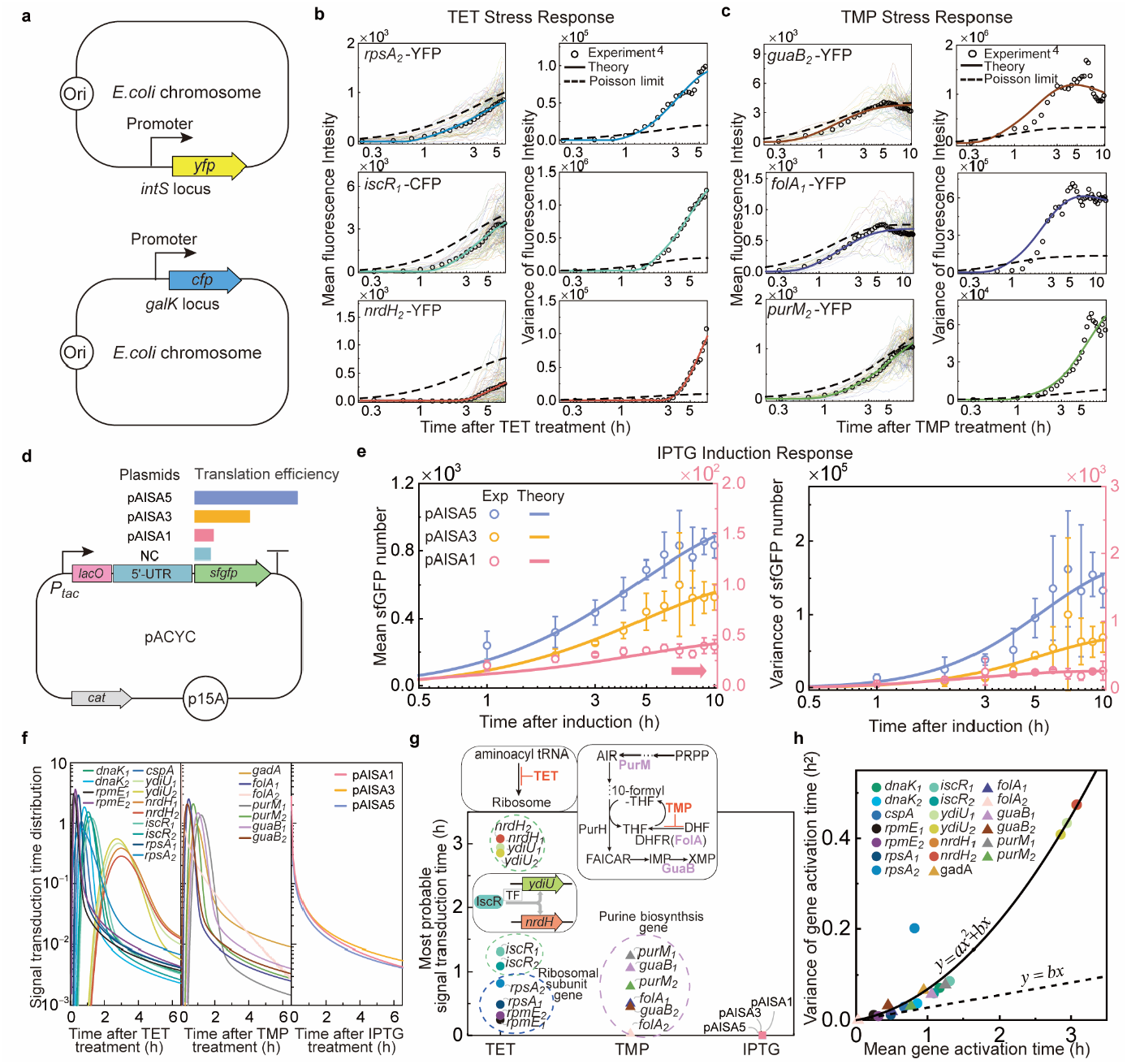
Quantitative analysis of adaptive gene expression dynamics under antibiotic stress and transcriptional induction. **a**, Schematic of chromosomal integration of promoter-reporter gene constructs. **b–c**, Time-dependent mean and variance of reporter gene (*yfp* or *cfp*) expression levels in response to antibiotic stress: (**b**) TET and (**c**) TMP. Subscript labels attached to promoter names indicate one of two independent datasets collected from different microcolonies. The symbols and solid lines represent experimental results from single-cell gene expression time traces^4^ (faint lines behind the mean profiles) and best fits of equations (1a) and (1b), each modified by a number-to-fluorescence intensity conversion factor (see Supplementary Note 10 and 11 for details of the quantitative analysis; additional results are shown in Supplementary Fig. 2). The dashed lines indicate the Poisson limit, in which all non-exponential waiting time distributions determined from the quantitative analysis are replaced by exponential distributions with the same mean. **d**, Plasmid constructs designed for IPTG-induced *sfgfp* gene expression. Different plasmid constructs contain distinct 5′-UTR sequences to control translation efficiency (see Experimental Methods and Supplementary Tables 1 and 2 for more details). **e**, Time-dependent mean and variance of sfGFP copy numbers in response to IPTG induction. The symbols and lines represent experimental results, with error bars indicating standard deviations over three independent experiments, and best fits of equations (1a) and (1b). **f**, Signal transduction time distributions extracted from our quantitative analyses. **g**, Signal transduction times and their associated biological processes. The symbols represent the most probable signal transduction times obtained from **f.** The colored dashed lines group promoters according to gene function: ribosomal protein genes (blue), transcription factor-regulated stress response genes (green), and purine biosynthesis genes (purple). The upper diagram illustrates the mode of action for TET and TMP. **h**, Relationship between the mean and variance of gene activation times extracted from our quantitative analysis of the antibiotic stress response data. In contrast to the linear mean-variance relationship for the protein maturation time (Fig. 2c), the variance here exhibits a quadratic dependence on the mean, implying distinct multi-step sequential dynamics in which correlations between individual step completion times govern the observed mean-variance relationship (Supplementary Note 6). Optimized parameter values are summarized in Supplementary Tables 3 and 4.

When combined with our cell adaptation model, equations (1a) and (1b) provide a unified, quantitative explanation of all experimental datasets, 13 and 7 distinct sets obtained for *E. coli* under TET and TMP stress signals, respectively (Fig. 3b, c and see Supplementary Fig 2). In addition, using essentially the same model, equations (1a) and (1b) also quantitatively explain the three experimental datasets on the time-dependent mean and variance of the fluorescent protein levels expressed in response to the IPTG transcription-induction signals, for three *E. coli* plasmid systems with different translational efficacies, in a consistent manner (Fig. 3e). The maturation time distribution of YFP, CFP, and sfGFP, required for these quantitative analyses, were extracted from previously reported experimental data^43^ (see Supplementary Fig. 3). In these quantitative analyses, the fluorescent protein lifetime distribution is modeled as an exponential distribution (see Supplementary Fig. 4), and the burst size is modeled as a negative binomial distribution^44,45^.

The signal transduction time distribution extracted from our quantitative analysis exhibited distinct shape depending on external signal type. The extracted signal transduction time distribution is found to be a unimodal function, represented by the convolution of a sub-Poisson gamma and a super-Poisson gamma distribution, for both TET and TMP antibiotic stresses (Fig. 3f). This result indicates that the signal transduction of the antibiotic stresses are significantly contributed from sub-Poisson and super-Poisson processes in a sequential manner, which may be assigned to the multi-step activation of chromosomal *E. coli* genes in response to stress signals and the cell-state dependent first expression burst of the activated gene, respectively. In contrast, for the IPTG transcriptional induction signal, the signal transduction time is found to follow a monotonically decreasing, super-Poisson gamma distribution (Fig. 3f). This result follows when the gene activation occurs on a time scale far shorter than the super-Poisson, first gene expression burst process and is consistent with the fact that IPTG rapidly activates of the associated *tac* promoter by immediate inactivation of the *lac* repressor^46,47^.

The antimicrobial-stress signal-induced gene-activation time exhibits significant gene-to-gene variation. Under TET stress, which suppresses protein synthesis by inhibiting aminoacyl t-RNA binding to ribosomes, ribosomal subunit genes, such as *rpmE* and *rpsA*, required for ribosome synthesis, show the shortest gene-activation times (Fig. 3g). By contrast, *iscR* gene, which is activated in response to oxidative stress induced by the protein-synthesis inhibition under TET stress, has a longer gene-activation time than the ribosomal subunit genes but a shorter one than further downstream genes such as *ydiU* and *nrdH*, for which IscR serves as a transcription factor. Our quantitative analysis also confirms that downstream genes have longer gene activation times for *E. coli* genes activated in response to TMP stress, which inhibits purine biosynthesis. The stress signal-transduction time as well exhibits significant gene-to-gene variation (Fig. 3f).

On the other hand, the distributions of the IPTG signal-transduction time extracted from our experiments exhibit little variation across plasmid constructs with different 5′-UTRs. This result is consistent with the fact that these plasmids differ in translational efficiency but share the same IPTG signal-transduction process (Fig. 3f, g).

Our analysis reveals that a quantitative relationship exists between the mean and variance of gene-activation times, which holds across diverse stress-response genes in *E. coli*. Specifically, the variance of the gene-activation time is found to be a quadratic function of the mean activation time for most *E. coli* genes activated by TET and TMP stress (Fig. 3h). This quadratic relationship results when the reaction times of elementary reactions constituting the gene-activation process are positively correlated (see Supplementary Note 6). Such positive correlations are frequently observed among intracellular reaction processes, owing to the influence of the common cell environment on their reaction times^48-50^ (see Supplementary Note 6).

## Discussion

Key features of our chemical-dynamics model include its modular representation of the cellular reaction network and characterization of the chemical dynamics of reaction-network modules using RTDs. As demonstrated in this work, the RTD of a network module can be expressed in terms of the RTDs of its constituent subnetwork modules. If necessary, each subnetwork module can be further decomposed into even smaller networks, when more detailed mechanistic information is available. For example, in this study, the mature-protein creation network is decomposed into several subnetwork modules, including signal sensing and propagation leading to gene activation, gene-expression, and protein maturation. The gene expression module is further decomposed into the first and subsequent gene expression burst modules. This decomposition of the mature-protein creation network enables us to define the signal transduction time as the sum of gene-activation and the ensuing first gene-expression burst times and investigate the separate effects of signal transduction and protein maturation processes on the dynamics of mature-protein number. Given that experimental data for the chemical dynamics of submodules are available, we can construct more detailed models of the submodules. Such hierarchical network modeling allows a systematic development of intracellular reaction-network models with the desired level of detail and accuracy. Together with modern high-throughput experiments on cell dynamics and machine-learning techniques, the present hierarchical network modeling framework suggests a new direction for the systematic development of digital twins of living cells, which enable quantitative predictions of stochastic cell dynamics in response to various external stimuli.

The gene activation and fluorescent protein maturation are both multi-step processes, but their RTDs exhibit qualitatively different mean–variance relationships (Fig. 2c and 3h), reflecting their distinct physical origins. A cellular reaction network module has a RTD with the quadratic mean-variance relationship when the module comprises sequential multi-step reactions with positively correlated reaction times. This positive correlation originated from the coupling of the reaction times to common cell environments. This is the case for gene activation module (see Supplementary Note 6). On the other hand, the fluorescent protein maturation time has a linear mean-variance relationship, which signifies that the unimolecular elementary reactions composing the protein maturation are largely independent processes

Our analyses confirm that the nascent protein burst size increases with transcriptional efficiency, which ranges from approximately ten to over a hundred protein copies per burst (see Supplementary Fig. 5). In addition, the plasmid constructs are found to have much larger protein burst sizes than chromosomal *E. coli* genes, which we attribute to the higher gene copy number in the plasmid expression system^51^.

A comparison between the present and previous theories^12,52^ is provided in Supplementary Note 7 for lack of space.

### Theoretical Methods

The derivation of equations. (1a)-(1b) for the mean and variance of mature protein number is as follows. The creation of mature protein consists of two modules: immature protein creation and maturation. Let 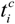 denote the time at which the *i*-th creation burst of immature proteins occurs and 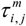 denotes the time required for the *j*-th immature protein created in the *i*-th burst to complete its maturation. Finally, 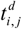 denotes the time at which the *j*-th immature protein created in the *i*-th burst is annihilated. Then, the number *n*(*t*) of mature protein at time is given by

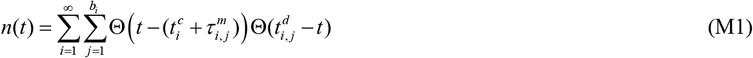

where *b*_*i*_ and Θ(*x*) denote, respectively, the number of immature proteins produced in the *i*-th burst and the Heaviside step function, which assumes 0 for negative *x* but 1 for positive *x* . We assume that 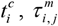 and *b*_*i*_ are independent random variables. Starting from equation M1, we obtain the analytical expressions of the mean and variance of mature protein numbers presented in equations. (1a)–(1b) (see Supplementary Method), without any further assumption.

In equation (1a), the mean creation rate of immature proteins is given by ⟨*R*_*T*_ (*t*) ⟩ = ⟨*b*⟩ ⟨*R*(*t*) ⟩ with ⟨*b*⟩ and ⟨*R*(*t*) ⟩ denoting the mean number of immature proteins per burst and the mean burst rate of immature proteins (see Supplementary Method). ⟨*R*(*t*) ⟩ is related to the signal transduction time distribution, *φ*_*st*_ (*t*) and the distribution of time interval between successive immature protein bursts, *φ*_*b*_ (*t*) . In the Laplace domain, this relationship simply reads as

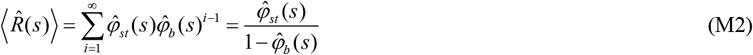

where 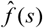 denotes the Laplace transform, i.e., 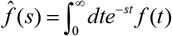. As shown in Fig. 1a, the signal transduction time is the sum of the target gene activation time, *t*_*ga*_, and the first gene expression burst time, 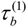, of the activated target gene. Then, *φ*_*st*_ (*t*) is given by the convolution of the distribution *φ*_*ga*_ (*t*) of gene activation time *t*_*ga*_ and the distribution 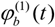 of the first gene expression burst time, 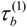, whose Laplace transform is given by

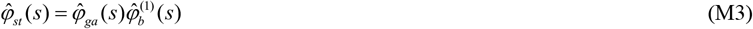

The Laplace transform of equations (1a) and (2) with (M2) is given by

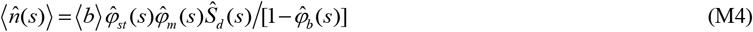

Equation (M4) shows that the mean number of mature proteins depends on the convolution of *φ*_*st*_ (*t*) and *φ*_*m*_ (*t*) .

In equation (1b), the TCF of fluctuations in the immature protein creation rate is given by

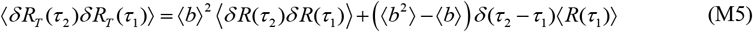

where ⟨ *b*^2^ ⟩ and ⟨ *δ R*(*τ*_2_)*δ R*(*τ*_1_) ⟩ denote, respectively, the mean square burst size and the TCF of fluctuations in the rate of the immature protein burst events. For our model, ⟨ *δ R*(*τ*_2_)*δ R*(*τ*_1_) ⟩ is related to the mean burst rate of immature proteins as

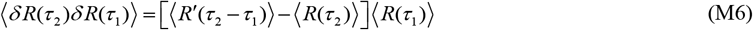

where ⟨ *R*′(*t*) ⟩ denotes the mean rate of the immature protein bursts following the first one. The Laplace domain expression of ⟨ *R*′(*t*) ⟩ is given by

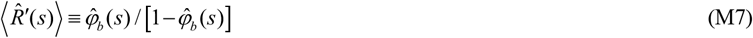

A more detailed derivations of these results are presented in Supplementary Note 4.

The steady-state mean of the mature protein number are obtained by applying Tauberian theorem to equation (M4), i.e.,

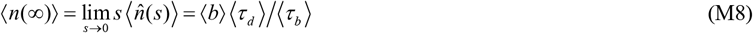

where ⟨*τ*_*d*_⟩ and ⟨ *τ*_*b*_ ⟩ denotes the mean lifetime of mature proteins and the mean interval of immature protein bursts. Equation (M8) follows from equation (M4) because 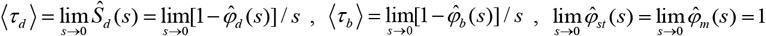. The steady-state variance of the mature protein number can be obtained from the long-time limit of equation (1b) (see Supplementary Note 5).

### Experimental Methods

#### Strains, plasmids, oligonucleotides, and reagents

All strains and plasmids used in this study are listed in Supplementary Table 1. *Escherichia coli* Mach1-T1^R^ (Invitrogen, Carlsbad, CA, USA) was used as the host strain for plasmid construction. The plasmids were prepared using a BIO4U^R^ AQprep™ Plasmid Extraction Kit (BIO4U, Seoul, Korea). DNA purification was performed using a GeneAll^R^ Expin™ CleanUp SV kit (GeneAll, Seoul, Korea) and Expin™ Gel SV kit (GeneAll, Seoul, Korea). All plasmid variants were assembled using the NEBuilder® HiFi DNA Assembly Kit (New England Biolabs, MA, USA). The Tac promoter and the terminator (BBa_B1006) were obtained from the Registry of Standard Biological Parts (https://parts.igem.org). Oligonucleotides for cloning were synthesized by Cosmogenetech (Seoul, Korea), and their sequences are listed in Supplementary Table 2. All reagents not specifically mentioned were purchased from Sigma-Aldrich (St. Louis, MO, USA).

A gene expression cassette with defined transcriptional strength (pAISA0) was constructed by assembling a synthetic promoter-*gfp* fragment with the pACYCDuet-1 as the backbone vector. The backbone vector was amplified using the primer pair pACYC_F/R and synthetic promoter-*gfp* contained Tac promoters were synthesized by Thermo Fisher Scientific (Waltham, MA, USA).

Following the construction of promoter variants with different transcriptional strengths, translation levels were further tuned by modulating the 5′-untranslated regions (5’-UTRs). Three 5′-UTR variants with distinct translational strengths were designed using the UTR Library Designer (https://sbi.postech.ac.kr/utr_library/)^41,42^ and introduced into the promoter constructs (pAISA0) to generate 3 combinatorial gene expression cassettes (pAISA1, pAISA3 and pAISA5) using the primer pairs UTR1_Tac_F/Tac_R, UTR3_Tac_F/Tac_R, UTR5_Tac_F/Tac_R.

#### sfGFP purification

The pQE80L_sfgfp was constructed by Gibson assembly of the pQE80L-mK and pACS00 fragments, which were PCR-amplified using primer pairs pQE_F/pQE_R and sfgfp_F/sfgfp_R, respectively. The pQE80L_sfgfp was transformed into *E. coli* DH10B for *sfgfp* expression. Cells were cultivated in LB medium at 37 °C until the OD_600_ reached 0.5, and then expression of *sfgfp* was induced by the addition of 1 mM isopropyl β-D-1-thiogalactopyranoside (IPTG). Following 6 hours of incubation, the cells were harvested by centrifugation at 13000 rpm and 4 °C for 15 min, and the cell pellets were stored at -80 °C until further use. The N-terminal

His6-tagged sgfp was purified using a MagListo His-tagged Protein Purification Kit (Bioneer, Korea) according to the manufacturer’s protocol. The purified protein was buffer-exchanged into 10 mM phosphate buffer (pH 7.4) using a centrifugal ultrafiltration filter (MWCO: 10,000; Merck Millipore, Billerica, MA, USA) and stored in 50% (v/v) glycerol.

#### Bacterial Culture and Preparation of Experimental Samples

Bacterial stocks stored at −80 °C were lightly touched with a pipette tip and inoculated into 3 mL of fresh LB medium supplemented with chloramphenicol (LB-Cm). Cultures were incubated overnight (∼12 h) at 37 °C with shaking at 200 rpm. A total of 0.15 mL of overnight culture was transferred into 2.85 mL of fresh LB-Cm and grown for ∼2 h to obtain exponentially growing cells. Cells were streaked onto LB-Cm agar plates and incubated for ∼10 h to generate colonies. Single colonies were used for all subsequent experiments. For time-course experiments, a single colony was inoculated into 3 mL of fresh LB-Cm medium and grown overnight (∼12 h). A total of 0.15 mL of the overnight culture was then diluted into 2.85 mL of fresh LB-Cm, which was designated as the 0 h time point for all experiments.

#### Absorbance Measurement

Optical density at 600 nm (OD_600_) was measured at designated time intervals to monitor bacterial growth. Fluorescence from sfGFP (Ex: 488 nm / Em: 510 nm) was measured to monitor response following transcription induction. Imaging and fluorescence measurements were restricted to the first 10 h to ensure that cells remained in exponential growth.

#### sfGFP Degradation Time Assay

A single colony was inoculated into 3 mL of LB medium and cultured overnight (∼12 h). The culture was diluted by transferring 0.15 mL into 2.85 mL of fresh LB containing 1 mM IPTG, followed by overnight induction (∼12 h). After induction, 1 mL of culture was collected and centrifuged at 5000 rpm for 5 min. The pellet was washed five times with PBS buffer to remove residual IPTG and resuspended in 1 mL of fresh LB without IPTG. A total of 0.15 mL of the washed culture was transferred into 2.85 mL of fresh LB, and fluorescence decay was monitored during incubation.

#### Image Intensity Correction

Non-uniform illumination resulted in higher intensities near the center of the image and reduced intensities toward the edges, introducing artificial variation in measured fluorescence levels. To correct for this effect, an even layer of purified sfGFP was applied to the imaging dish, and the illumination profile was recorded under identical imaging conditions. Region-of-interest (ROI) masks corresponding to each imaging position were applied to compute local correction factors. Corrected fluorescence values were rescaled to match the dynamic range of raw measurements. This correction substantially reduced abrupt fluctuations in fluorescence intensity, confirming that the observed variability primarily arose from uneven illumination rather than biological heterogeneity.

#### Conversion of fluorescence intensity to protein copy number

Bacterial cells in a target sample were imaged, and the integrated fluorescence intensity for each cellular ROI was measured. Background correction was performed by subtracting the integrated intensity measured for the same ROI in a background sample under identical optical and acquisition settings. To convert the resulting fluorescence intensities into absolute protein copy numbers, a reference sample containing 1 μM purified sfGFP was imaged under the same conditions. Within this reference sample, integrated fluorescence intensities were measured from ROIs, each representing a specific cellular ROI, and background correction was applied as above. The detection volume for each ROI was estimated as the product of the ROI area and the axial detection depth. Using the known sfGFP concentration, the total number of sfGFP molecules within this volume was calculated, which allows derivation of a conversion factor corresponding to the background-corrected fluorescence intensity per single sfGFP molecule. Finally, the absolute protein copy number per cell in the target sample was obtained by dividing the background-corrected integrated fluorescence intensity of each cell by this conversion factor.

## Data availability

The time-lapse microscopy data used in Figs. 3b,c and Supplementary Fig. 2 are publicly available from ref. 4. The data used in Fig. 3e are provided in the Source Data file. The data analyzed in Supplementary Fig. 3 were obtained from the previously published dataset described in ref. 43.

## Code availability

All code for data analysis and stochastic simulation is available on (https://github.com/Jinhyung96/cellular-chemical-dynamics)

## Acknowledgements

This work was supported by Global Science Research Center Program (RS-2024-00411134) (J.S., J.-H.K. and H.R.K.). and Chung-Ang University Graduate Research Scholarship in 2021.

## Author contributions

J.K., S.J.P. and J.S. derived the theoretical equations. J.K. and J.-H.K. performed the data analysis; J.K. performed the stochastic simulations. S.K., K.J., G.Y.J., S.J. and H.R.K. performed the transcriptional induction experiments. J.K., J.-H.K., K.J., S.S., G.Y.J., S.J. and J.S. wrote the manuscript. J.-H.K. and J.S. designed the research. J.S. supervised the project.

