## Supplementary Information for "Cellular Chemical Dynamics Governing Signal Transduction and Adaptive Gene Expression: Beyond Classical Kinetics"

### SUPPLEMENTARY METHODS

In this section, we present a derivation of equations (1a) and (1b) for the general cell adaptation model present in the main text. These equations describe time dependence of the mean and variance of the number of mature proteins undergoing creation and annihilation processes. The creation of mature protein involves two sequential steps: the creation of immature proteins and the subsequent maturation. We assume that multiple immature proteins are created at a single gene expression burst. Immature proteins thus created undergo maturation. During the maturation process, new bursts of immature proteins occur. Let $t_i^c$ denote the time at which the *i*-th creation burst of immature proteins occurs and $\tau_{i,j}^m$ denotes the time required for the *j*-th immature protein created in the *i*-th burst to complete its maturation. Similarly, $t_{i,j}^d$ denotes the time at which the *j*-th immature protein created in the *i*-th burst is annihilated. Then, the number *n*(t) of mature protein at time is given by equation (M1) in the Methods section of the main text.

**The mean of the mature protein number**

Equation (M1) can be rewritten as

$$n(t) = \sum_{i=1}^{\infty}\sum_{j=1}^{b_i} \Theta\left(t - t_i^c\right)\Theta\left(t - (t_i^c + \tau_{i,j}^m)\right)\Theta(t_{i,j}^d - t) \tag{M1-1}$$

where we have used the following identity, $\Theta\left(t-(t_i^c+\tau_{i,j}^m)\right) = \Theta\left(t-t_i^c\right)\Theta\left(t-(t_i^c+\tau_{i,j}^m)\right)$, which holds for Heavyside step function $\Theta$ as long as $\tau_{i,j}^m > 0$. In equation (M1-1), $\Theta(t_{i,j}^d - t)$ can be replaced by $1-\Theta(t-t_{i,j}^d)$. Using

$\Theta(t-x)=\int_0^t d\tau\delta(\tau-x)=\int_0^a d\tau\delta(\tau-x)+\int_a^t d\tau\delta(\tau-x)$, which holds for any constant $a$, we decompose $\Theta(t-t_{i,j}^d)$ into

$$\Theta(t-t_{i,j}^d)=\int_0^t d\tau\delta(\tau-t_{i,j}^d)=\int_0^{t_i^c+\tau_{i,j}^m} d\tau\delta(\tau-t_{i,j}^d)+\int_{t_i^c+\tau_{i,j}^m}^t d\tau\delta(\tau-t_{i,j}^d) \quad \text{(M1-2)}$$

where $\delta(x)$ denotes the Dirac delta function. Because the mature protein degradation time, $t_{i,j}^d$, is always greater than the mature protein creation time, $t_i^c+\tau_{i,j}^m$, the first integral on the right-hand side of equation (M1-2) vanishes, i.e., $\int_0^{t_i^c+\tau_{i,j}^m} d\tau\delta(\tau-t_{i,j}^d)=0$. By changing the integration variable from $\tau$ to $\tau'=\tau-(t_i^c+\tau_{i,j}^m)$ in the second integral on the right-hand-side of equation (M1-2), we obtain

$$\Theta(t-t_{i,j}^d)=\int_0^{t-(t_i^c+\tau_{i,j}^m)} d\tau'\delta\left(\tau'-\left(t_{i,j}^d-(t_i^c+\tau_{i,j}^m)\right)\right)=\int_0^{t-(t_i^c+\tau_{i,j}^m)} d\tau'\delta\left(\tau'-\tau_{i,j}^d\right) \quad \text{(M1-3)}$$

where $\tau_{i,j}^d\left(=t_{i,j}^d-t_i^c-\tau_{i,j}^m\right)$ denotes the lifetime of the mature protein. Using equation (M1-3), we can rewrite equation (M1-1) as

$$\begin{aligned}
n(t)&=\sum_{i=1}^{\infty}\sum_{j=1}^{b_i}\int_0^t d\tau\delta(\tau-t_i^c)\Theta(t-t_i^c-\tau_{i,j}^m)\left(1-\int_0^{t-t_i^c-\tau_{i,j}^m} d\tau'\delta(\tau'-\tau_{i,j}^d)\right)\\
&=\sum_{i=1}^{\infty}\sum_{j=1}^{b_i}\int_0^t d\tau\delta(\tau-t_i^c)\int_0^{t-t_i^c} dy\delta(y-\tau_{i,j}^m)\left(1-\int_0^{t-t_i^c-\tau_{i,j}^m} d\tau'\delta(\tau'-\tau_{i,j}^d)\right) \quad \text{(M1-4)}\\
&=\sum_{i=1}^{\infty}\sum_{j=1}^{b_i}\int_0^t d\tau\delta(\tau-t_i^c)\int_0^{t-\tau} dy\delta(y-\tau_{i,j}^m)\left(1-\int_0^{t-\tau-y} d\tau'\delta(\tau'-\tau_{i,j}^d)\right)
\end{aligned}$$

The last equation in equation (M1-4) follows from the second because $\delta(y-\tau_{i,j}^m)f(\tau_{i,j}^m)=\delta(y-\tau_{i,j}^m)f(y)$ for any function $f$. By taking the average of equation (M1-4) over distributions of $\{b_i,t_i^c,\tau_{i,j}^m,\tau_{i,j}^d\}$, we obtain

$$\langle n(t)\rangle = \langle b\rangle \sum_{i=1}^{\infty} \int_0^t d\tau \langle \delta(\tau - t_i^c)\rangle \int_0^{t-\tau} dy \langle \delta(y - \tau_{i,j}^m)\rangle \left(1 - \int_0^{t-\tau-y} d\tau' \langle \delta(\tau' - \tau_{i,j}^d)\rangle\right) \quad \text{(M1-5)}$$

where we have used the law of the total expectation, i.e., $\left\langle \sum_{j=1}^{b_i} f_j \right\rangle = \langle b_i \rangle \langle f_j \rangle$ and our assumption that the mean burst size is the same for all bursts, $\langle b_i \rangle = \langle b \rangle$.

$\langle \delta(y - \tau_{i,j}^m)\rangle$ and $\langle \delta(\tau' - \tau_{i,j}^d)\rangle$ appearing in equation (M1-5) are the probability distribution $\varphi_{i,j}^m(t)$ of maturation time $\tau_{i,j}^m$ of immature protein and the probability distribution $\varphi_{i,j}^d(t)$ of lifetime $\tau_{i,j}^d$ of mature protein. This follows because $\langle \delta(\tau - x)\rangle = \int_0^{\infty} \delta(\tau - x) f(x) dx = f(\tau)$ for any continuous random variable $x$ with probability distribution $f(x)$. Then, equation (M1-5) can be written in terms of $\varphi_{i,j}^m(t)$ and $\varphi_{i,j}^d(t)$ as follows:

$$\langle n(t)\rangle = \langle b\rangle \sum_{i=1}^{\infty} \int_0^t d\tau \langle \delta(\tau - t_i^c)\rangle \int_0^{t-\tau} dy \varphi_{i,j}^m(y) \left(1 - \int_0^{t-\tau-y} d\tau' \varphi_{i,j}^d(\tau')\right) \quad \text{(M1-6)}$$

Here, $\varphi_{i,j}^m(t)$ denotes the distribution of maturation time of the *j*-th nascent protein produced in the *i*-th burst. $\varphi_{i,j}^d(t)$ denotes the lifetime distribution of the corresponding mature protein. Given that all mature proteins share the same lifetime distribution and maturation time distribution, i.e., $\varphi_{i,j}^d(t) = \varphi_d(t)$ and $\varphi_{i,j}^m(t) = \varphi_m(t)$ for any *i*, *j*, we can rewrite equation (M1-6) as

$$\langle n(t)\rangle = \langle b\rangle \int_0^t d\tau \left\langle \sum_{i=1}^{\infty} \delta(\tau - t_i^c) \right\rangle P_m(t - \tau) \quad \text{(M1-7)}$$

where $P_m(t)=\int_0^t dy\varphi_m(y)\left(1-\int_0^{t-y}dz\varphi_d(z)\right)$, which is the convolution of $\varphi_m(t)$ and the survival probability of the mature protein $S_d(t)\equiv 1-\int_0^t d\tau\varphi_d(\tau)=\int_t^\infty d\tau\varphi_d(\tau)$. $P_m(t)$ designates the probability that an immature protein created at time 0 completes maturation before time *t* and survives until time *t*. This expression is identical to equation (2) in the main text. We define the mean burst rate of immature proteins by $\langle R(t)\rangle=\sum_{i=1}^{\infty}\langle\delta(\tau-t_i^c)\rangle$. By substituting $\langle R(t)\rangle=\sum_{i=1}^{\infty}\langle\delta(\tau-t_i^c)\rangle$ and using the relationship between the mean burst rate and the mean creation rate of immature proteins, i.e., $\langle R_T(\tau)\rangle\equiv\langle b\rangle\langle R(\tau)\rangle$, into equation (M1-7), we obtain equation (1a) in the main text.

#### The variance of the mature protein number

The derivation of equation (1b) is similar to the derivation of equation (1a). From equation (M1-1), we obtain the following equation for $n^2(t)$:

$$
\begin{aligned}
n^2(t) &= \left(\sum_{i=1}^{\infty}\Theta\left(t-t_i^c\right)\sum_{j=1}^{b_i}\Theta\left(t-t_i^c-\tau_{i,j}^m\right)\left(1-\Theta\left(t-t_{i,j}^d\right)\right)\right)^2 \\
&= \sum_{i=1}^{\infty}\Theta\left(t-t_i^c\right)\left(\sum_{j=1}^{b_i}\Theta\left(t-t_i^c-\tau_{i,j}^m\right)\left(1-\Theta\left(t-t_{i,j}^d\right)\right)\right)^2 \\
&+ \sum_{i=1}^{\infty}\sum_{\substack{k=1\\ i\neq k}}^{\infty}\Theta\left(t-t_i^c\right)\Theta\left(t-t_k^c\right)\sum_{j=1}^{b_i}\Theta\left(t-t_i^c-\tau_{i,j}^m\right)\left(1-\Theta\left(t-t_{i,j}^d\right)\right)\sum_{l=1}^{b_k}\Theta\left(t-t_k^c-t_{k,l}^m\right)\left(1-\Theta\left(t-t_{k,l}^d\right)\right)
\end{aligned}
$$

(M1-8)

where we have used $\Theta^2(t-a)=\Theta(t-a)$ in obtaining the second equality.

By taking the average of $n^2(t)$ over $\{t_i^c, t_k^c\}$, $\{\tau_{i,j}^m, \tau_{k,l}^m\}$, $\{t_{i,j}^d, t_{k,l}^d\}$, $\{b_i, b_k\}$, we obtain

$$\begin{aligned}\left\langle n(t)^2 \right\rangle &= \sum_{i=1}^{\infty} \left\langle \Theta\left(t-t_i^c\right)\right\rangle \left\langle \left( \sum_{j=1}^{b_i} \Theta\left(t-t_i^c-\tau_{i,j}^m\right)\left(1-\Theta\left(t-t_{i,j}^d\right)\right)\right)^2 \right\rangle \\ &+ \left\langle \sum_{i=1}^{\infty}\sum_{\substack{k=1\\ i\neq k}}^{\infty} \Theta\left(t-t_i^c\right)\Theta\left(t-t_k^c\right)\sum_{j=1}^{b_i}\Theta\left(t-t_i^c-\tau_{i,j}^m\right)\left(1-\Theta\left(t-t_{i,j}^d\right)\right)\sum_{l=1}^{b_k}\Theta\left(t-t_k^c-\tau_{k,l}^m\right)\left(1-\Theta\left(t-t_{k,l}^d\right)\right)\right\rangle\end{aligned}$$

(M1-9)

Given that $b_i$, $t_i^c$, $\tau_{i,j}^m$ and $t_{i,j}^d$ are independent random variables, we can obtain an analytic expression for $\left\langle \left( \sum_{j=1}^{b_i} \Theta\left(t-t_i^c-\tau_{i,j}^m\right)\left(1-\Theta\left(t-t_{i,j}^d\right)\right)\right)^2 \right\rangle$ in equation (M1-9) by using the following identity,

$$\begin{aligned}\left\langle \left(\sum_{j=1}^{N} a_j\right)^2 \right\rangle &= \left\langle \delta a^2 \right\rangle \left\langle N \right\rangle + \left\langle N^2 \right\rangle \left\langle a \right\rangle^2 \\ &= \left[\left\langle a^2 \right\rangle - \left\langle a \right\rangle^2\right]\left\langle N \right\rangle + \left\langle N^2 \right\rangle \left\langle a \right\rangle^2\end{aligned},$$

where $\left\langle \delta a^2 \right\rangle$ denotes the variance of $a_i$. This identity can be obtained by combining the law of the total expectation, i.e. $\left\langle \left(\sum_{j=1}^{N} a_j\right) \right\rangle = \left\langle N \right\rangle \left\langle a \right\rangle$, and the law of total variance, i.e. $\left\langle \delta\left(\sum_{j=1}^{N} a_j\right)^2 \right\rangle = \left\langle \left(\sum_{j=1}^{N} a_j\right)^2 \right\rangle - \left\langle \left(\sum_{j=1}^{N} a_j\right) \right\rangle^2 = \left\langle \delta a^2 \right\rangle \left\langle N \right\rangle + \left\langle \delta N^2 \right\rangle \left\langle a \right\rangle^2$, which holds whenever $a_i$ and $N$ are independent random variables. Here, $\left\langle \delta N^2 \right\rangle$ denotes the variances of $N$. By identifying $a_j = \Theta\left(t-t_i^c-\tau_{i,j}^m\right)\left(1-\Theta\left(t-t_{i,j}^d\right)\right)$ and $N = b_i$, we obtain the following expression for $\left\langle \left( \sum_{j=1}^{b_i} \Theta\left(t - t_i^c - \tau_{i,j}^m\right)\left(1 - \Theta\left(t - t_{i,j}^d\right)\right) \right)^2 \right\rangle$

$$\begin{aligned}
&\left\langle \left( \sum_{j=1}^{b_i} \Theta\left(t - t_i^c - \tau_{i,j}^m\right)\left(1 - \Theta\left(t - t_{i,j}^d\right)\right) \right)^2 \right\rangle \\
&= \left\langle \delta\left( \Theta\left(t - t_i^c - \tau_{i,j}^m\right)\left(1 - \Theta\left(t - t_{i,j}^d\right)\right) \right)^2 \right\rangle \langle b \rangle + \left\langle b^2 \right\rangle \left\langle \Theta\left(t - t_i^c - \tau_{i,j}^m\right)\left(1 - \Theta\left(t - t_{i,j}^d\right)\right) \right\rangle^2 \\
&= \left( \left\langle \left( \Theta\left(t - t_i^c - \tau_{i,j}^m\right)\left(1 - \Theta\left(t - t_{i,j}^d\right)\right) \right)^2 \right\rangle - \left\langle \Theta\left(t - t_i^c - \tau_{i,j}^m\right)\left(1 - \Theta\left(t - t_{i,j}^d\right)\right) \right\rangle^2 \right) \langle b \rangle \\
&\quad + \left\langle b^2 \right\rangle \left\langle \Theta\left(t - t_i^c - \tau_{i,j}^m\right)\left(1 - \Theta\left(t - t_{i,j}^d\right)\right) \right\rangle^2 \\
&= \left\langle \left( \Theta\left(t - t_i^c - \tau_{i,j}^m\right)\left(1 - \Theta\left(t - t_{i,j}^d\right)\right) \right)^2 \right\rangle \langle b \rangle + \left( \left\langle b^2 \right\rangle - \langle b \rangle \right) \left\langle \Theta\left(t - t_i^c - \tau_{i,j}^m\right)\left(1 - \Theta\left(t - t_{i,j}^d\right)\right) \right\rangle^2
\end{aligned}$$

(M1-10)

where $\langle b^n \rangle$ denotes the $n$-th moment of the burst size distribution. Using the identity $\Theta(x)^2 = \Theta(x)$, we can rewrite $\left\langle \left( \Theta\left(t - t_i^c - \tau_{i,j}^m\right)\left(1 - \Theta\left(t - t_{i,j}^d\right)\right) \right)^2 \right\rangle$ in the first term on the right-hand-side of the last equality in equation (M1-10) as follows:

$$\begin{aligned}
&\left\langle \left( \Theta\left(t - t_i^c - \tau_{i,j}^m\right)\left(1 - \Theta\left(t - t_{i,j}^d\right)\right) \right)^2 \right\rangle \\
&= \left\langle \Theta\left(t - t_i^c - \tau_{i,j}^m\right)^2 \left(1 - 2\Theta\left(t - t_{i,j}^d\right) + \Theta\left(t - t_{i,j}^d\right)^2 \right) \right\rangle \\
&= \left\langle \Theta\left(t - t_i^c - \tau_{i,j}^m\right)\left(1 - \Theta\left(t - t_{i,j}^d\right)\right) \right\rangle
\end{aligned} \qquad \text{(M1-11)}$$

Substituting equation (M1-11) into the right-hand-side of the last equality in equation (M1-10) we obtain

$$\left\langle \left( \sum_{j=1}^{b_i} \Theta\left(t - t_i^c - \tau_{i,j}^m\right)\left(1 - \Theta\left(t - t_{i,j}^d\right)\right) \right)^2 \right\rangle$$

$$= \left\langle \Theta\left(t - t_i^c - \tau_{i,j}^m\right)\left(1 - \Theta\left(t - t_{i,j}^d\right)\right) \right\rangle \langle b \rangle + \left( \langle b^2 \rangle - \langle b \rangle \right) \left\langle \Theta\left(t - t_i^c - \tau_{i,j}^m\right)\left(1 - \Theta\left(t - t_{i,j}^d\right)\right) \right\rangle^2$$

(M1-12)

By substituting equation (M1-12) into the right-hand-side of equation (M1-9) and using identity $\Theta(x)^2 = \Theta(x)$, we obtain

$$\left\langle n(t)^2 \right\rangle = \sum_{i=1}^{\infty} \left\langle \Theta\left(t - t_i^c\right) \right\rangle \begin{pmatrix} \left\langle \Theta\left(t - t_i^c - \tau_{i,j}^m\right)\left(1 - \Theta\left(t - t_{i,j}^d\right)\right) \right\rangle \langle b \rangle \\ + \left( \langle b^2 \rangle - \langle b \rangle \right) \left\langle \Theta\left(t - t_i^c - \tau_{i,j}^m\right)\left(1 - \Theta\left(t - t_{i,j}^d\right)\right) \right\rangle^2 \end{pmatrix}$$
$$+ \left\langle \sum_{i=1}^{\infty} \sum_{\substack{k=1 \\ i \neq k}}^{\infty} \Theta\left(t - t_i^c\right) \Theta\left(t - t_k^c\right) \sum_{j=1}^{b_i} \Theta\left(t - t_i^c - \tau_{i,j}^m\right)\left(1 - \Theta\left(t - t_{i,j}^d\right)\right) \sum_{l=1}^{b_k} \Theta\left(t - t_k^c - \tau_{k,l}^m\right)\left(1 - \Theta\left(t - t_{k,l}^d\right)\right) \right\rangle$$

(M1-13)

Using the same method employed to derive equation (M1-4) from equation (M1-1), we obtain the following equation from equation (M1-13)

$$\left\langle n(t)^2 \right\rangle = \langle b \rangle \sum_{i=1}^{\infty} \int_0^t d\tau \langle \delta(\tau - t_i^c) \rangle \int_0^{t-\tau} dy \langle \delta(y - \tau_{i,j}^m) \rangle \left( 1 - \int_0^{t-\tau-y} d\tau' \langle \delta(\tau' - \tau_{i,j}^d) \rangle \right)$$
$$+ \left( \langle b^2 \rangle - \langle b \rangle \right) \sum_{i=1}^{\infty} \int_0^t d\tau \langle \delta(\tau - t_i^c) \rangle \left( \int_0^{t-\tau} dy \langle \delta(y - \tau_{i,j}^m) \rangle \left( 1 - \int_0^{t-\tau-y} d\tau' \langle \delta(\tau' - \tau_{i,j}^d) \rangle \right) \right)^2$$
$$+ \langle b \rangle^2 \sum_{i=1}^{\infty} \sum_{\substack{k=1 \\ i \neq k}}^{\infty} \int_0^t d\tau_1 \int_0^t d\tau_2 \langle \delta(\tau_1 - t_i^c) \delta(\tau_2 - t_k^c) \rangle \begin{pmatrix} \int_0^{t-\tau_1} dy \langle \delta(y - \tau_{i,j}^m) \rangle \left( 1 - \int_0^{t-\tau_1-y} d\tau_1' \langle \delta(\tau_1' - \tau_{i,j}^d) \rangle \right) \\ \times \int_0^{t-\tau_2} dy \langle \delta(y - \tau_{k,l}^m) \rangle \left( 1 - \int_0^{t-\tau_2-y} d\tau_2' \langle \delta(\tau_2' - \tau_{k,l}^d) \rangle \right) \end{pmatrix}$$

(M1-14)

Given that all mature proteins share the same lifetime distribution, i.e., given that $\varphi_d(\tau) = \langle \delta(\tau - \tau_{i,j}^d) \rangle$ for all $i$ and $j$, and all immature proteins have the same maturation distribution, i.e., $\varphi_m(y) = \langle \delta(y - \tau_{i,j}^m) \rangle$ for all $i$ and $j$, we can rewrite equation (M1-14) as

$$\begin{aligned} \left\langle n(t)^2 \right\rangle &= \langle b \rangle \int_0^t d\tau \left\langle \sum_{i=1}^{\infty} \delta(\tau - t_i^c) \right\rangle P_m(t-\tau) \\ &+ \left( \left\langle b^2 \right\rangle - \langle b \rangle \right) \int_0^t d\tau \left\langle \sum_{i=1}^{\infty} \delta(\tau - t_i^c) \right\rangle P_m(t-\tau)^2 \\ &+ \langle b \rangle^2 \int_0^t d\tau_1 \int_0^t d\tau_2 \left\langle \sum_{i=1}^{\infty} \sum_{\substack{k=1 \\ i \neq k}}^{\infty} \delta(\tau_1 - t_i^c) \delta(\tau_2 - t_k^c) \right\rangle P_m(t-\tau_1) P_m(t-\tau_2) \end{aligned} \tag{M1-15}$$

where $P_m(t) = \int_0^t dy \varphi_m(y) \left(1 - \int_0^{t-y} dz \varphi_d(z)\right)$. The first term in equation (M1-15) is the same as $\langle n(t) \rangle$ given in equation (M1-7). $\left\langle \sum_{i=1}^{\infty} \sum_{\substack{k=1 \\ i \neq k}}^{\infty} \delta(\tau_1 - t_i^c) \delta(\tau_2 - t_k^c) \right\rangle$ in the last term on the right-hand-side of equation (M1-15) is defined as the time correlation function (TCF) of the immature protein burst events[1], i.e.,

$$\left\langle R(\tau_1) R(\tau_2) \right\rangle \equiv \left\langle \sum_{i=1}^{\infty} \sum_{\substack{k=1 \\ i \neq k}}^{\infty} \delta(\tau_1 - t_i^c) \delta(\tau_2 - t_k^c) \right\rangle \tag{M1-16}$$

Note that the diagonal terms with $i = j$ do not make any contribution to the TCF of the immature protein burst events, given in equation (M1-16). When immature protein burst event is renewal process, the TCF given in equation (M1-16) reduces to $\left\langle R(\tau_1) \right\rangle \left\langle R(\tau_2 - \tau_1) \right\rangle$ (see Supplementary Note 4). When the burst event is Poisson process, for which the mean burst rate is constant in time, i.e., $\left\langle R(\tau) \right\rangle = k$, the TCF of the burst rate fluctuation vanishes, i.e., $\left\langle \delta R(\tau_1) \delta R(\tau_2) \right\rangle = \left\langle R(\tau_1) R(\tau_2) \right\rangle - \left\langle R(\tau_1) \right\rangle \left\langle R(\tau_2) \right\rangle = 0$ because the TCF given in equation (M1-16) reduces to $\left\langle R(\tau_1) \right\rangle \left\langle R(\tau_2) \right\rangle = k^2$.

Substituting equation (M1-16) into equation (M1-13), we obtain

$$\begin{aligned}\left\langle n(t)^2\right\rangle &= \langle n(t)\rangle + \left(\left\langle b^2\right\rangle - \langle b\rangle\right)\int_0^t d\tau \left\langle R(\tau)\right\rangle P_m(t-\tau)^2 \\ &+ \langle b\rangle^2 \int_0^t d\tau_2 \int_0^t d\tau_1 \left\langle R(\tau_1)R(\tau_2)\right\rangle P_m(t-\tau_1)P_m(t-\tau_2)\end{aligned} \quad \text{(M1-17)}$$

Subtracting $\langle n(t)\rangle^2$ from equation (M1-17), we obtain the analytic expression for the variance $\sigma_n^2(t)\left(=\left\langle n(t)^2\right\rangle - \left\langle n(t)\right\rangle^2\right)$ of product number, equation (1b).

$$\begin{aligned}\sigma_n{}^2(t) &= \langle n(t)\rangle \\ &+ \left(\left\langle b^2\right\rangle - \langle b\rangle\right)\int_0^t d\tau \left\langle R(\tau)\right\rangle P_m(t-\tau)^2 \\ &+ \left\langle b\right\rangle^2 \int_0^t d\tau_2 \int_0^t d\tau_1 P_m(t-\tau_1)P_m(t-\tau_2)\left[\left\langle R(\tau_2)R(\tau_1)\right\rangle - \left\langle R(\tau_2)\right\rangle\left\langle R(\tau_1)\right\rangle\right]\end{aligned} \quad \text{(M1-18)}$$

In obtaining equation (M1-18), we have used the following identity:

$$\left\langle n(t)\right\rangle^2 = \langle b\rangle^2 \int_0^t d\tau_2 \int_0^t d\tau_1 \left\langle R(\tau_2)\right\rangle\left\langle R(\tau_1)\right\rangle P_m(t-\tau_2)P_m(t-\tau_1) \quad \text{(M1-19)}$$

which follows from equation (1a) in the main text.

We obtain equation (1b) from equation (M1-18). The integral in the second term on the right-hand-side of equation (M1-18) can be written as

$$\int_0^t d\tau \left\langle R(\tau)\right\rangle P_m(t-\tau)^2 = \int_0^t d\tau \int_0^t dx\,\delta(\tau-x)\left\langle R(x)\right\rangle P_m(t-x)P_m(t-\tau) \quad \text{(M1-20)}$$

which results from the following property of Dirac's delta, $\int_0^t dx\,\delta(\tau-x)f(x) = f(\tau)$ given that $t>\tau$. Using this equality, we rewrite equation (M1-18) as

$$\sigma_n^{\,2}(t) = \langle n(t)\rangle + \left(\left\langle b^2\right\rangle - \langle b\rangle\right)\int_0^t d\tau_2 \int_0^t d\tau_1 \delta(\tau_2-\tau_1)\left\langle R(\tau_1)\right\rangle P_m(t-\tau_1)P_m(t-\tau_2) + \langle b\rangle^2 \int_0^t d\tau_2 \int_0^t d\tau_1 P_m(t-\tau_1)P_m(t-\tau_2)\left[\left\langle R(\tau_2)R(\tau_1)\right\rangle - \left\langle R(\tau_2)\right\rangle\left\langle R(\tau_1)\right\rangle\right] \tag{M1-21}$$

Equation (M1-21) yields equation (1b) in the main text, where the time correlation function $\langle \delta R_T(\tau_2)\delta R_T(\tau_1)\rangle$ of the immature protein creation rate fluctuation is defined as

$$\langle \delta R_T(\tau_2)\delta R_T(\tau_1)\rangle\left[= \langle b\rangle^2 \left\langle \delta R(\tau_2)\delta R(\tau_1)\right\rangle + \left(\langle b^2\rangle - \langle b\rangle\right)\delta(\tau_2-\tau_1)\langle R(\tau_1)\rangle\right]. \tag{M1-22}$$

Here $\langle \delta R(\tau_2)\delta R(\tau_1)\rangle$ denotes the TCF of immature protein burst events, defined by

$$\langle \delta R(\tau_2)\delta R(\tau_1)\rangle = \langle R(\tau_2)R(\tau_1)\rangle - \langle R(\tau_2)\rangle\langle R(\tau_1)\rangle \tag{M1-23}$$

When the burst size is unity, i.e., when $\left\langle b^2\right\rangle = \left\langle b\right\rangle = 1$, $\langle \delta R_T(\tau_2)\delta R_T(\tau_1)\rangle$ reduces to $\left\langle \delta R(\tau_2)\delta R(\tau_1)\right\rangle$. The functional form of $\left\langle \delta R(\tau_2)\delta R(\tau_1)\right\rangle$ can be related to the RTDs, $\varphi_{st}(t)$ and $\varphi_b(t)$ of the signal transduction and subsequent gene expression processes (Supplementary Note 4).

### SUPPLEMENTARY NOTES

#### Supplementary Note 1. Gamma Central Limit Theorem for Reaction Time Distribution of Arbitrary Multi-Step processes

In this note, we show that the reaction time distribution of a multi-step reaction process can be approximated by a gamma distribution. The total reaction time is defined by the sum of the reaction times, $\{\tau_i\}$, of individual elementary reactions, i.e., $T=\sum_{i=1}^{n}\tau_i$. $\{\tau_1,\tau_2,\cdots,\tau_n\}$ may follow different waiting time distributions and are assumed independent from each other. Then the Laplace transform of the distribution $\Phi(T)$ of $T$ is given by

$$\hat{\Phi}(s)=\prod_{i=1}^{n}\hat{\varphi}_i(s). \quad \text{(N1-1)}$$

where $\hat{\varphi}_i(s)$ denotes the Laplace transform of the distribution of $\tau_i$. In the small $s$ regime, $\hat{\varphi}_i(s)$ can be approximated by its Taylor series, i.e., $\hat{\varphi}_i(s)=1-\langle t_i\rangle s+\frac{1}{2}\langle t_i^2\rangle s^2+\cdots$.

Substituting this result into the logarithms of (N1-1), we obtain

$$\ln\hat{\Phi}(s)=\sum_{i=1}^{n}\ln\hat{\varphi}_i(s)=\sum_{i=1}^{n}\ln\left(1-\langle t_i\rangle s+\frac{1}{2}\langle t_i^2\rangle s^2+\cdots\right) \qquad (\langle t_i\rangle s<<1) \quad \text{(N1-2)}$$

Using $\ln(1+x)=x-\frac{x^2}{2}+\cdots$ $(x<<1)$, we obtain

$$\ln\hat{\Phi}(s)\cong\sum_{i=1}^{n}-\langle t_i\rangle s+\frac{1}{2}\left(\langle t_i^2\rangle-\langle t_i\rangle^2\right)s^2 \qquad (\langle t_i\rangle s<<1) \quad \text{(N1-3)}$$

This result can be rearranged to

$$\ln \hat{\Phi}(s) = -\langle T\rangle s + \frac{1}{2}\langle \delta T^2\rangle s^2 \tag{N1-4}$$

where $\langle T\rangle$ and $\langle \delta T^2\rangle$ are given by

$$\langle T\rangle = \sum_{i=1}^{n}\langle t_i\rangle \text{ and } \langle \delta T^2\rangle = \sum_{i=1}^{n}\langle \delta t_i^{\,2}\rangle . \tag{N1-5}$$

The distribution $\Phi(T)$ of total reaction time *T*, whose Laplace transform is given in equation (N1-4), can be approximated by a Gamma distribution with the mean and variance given by $\langle T\rangle$ and $\langle \delta T^2\rangle$. This can be easily proved by showing that the logarithm of the Laplace transform of a Gamma distribution approaches to the right-hand-side of equation (N1-4). The Laplace transform of a Gamma distribution $G(T)$ is given by

$$\hat{G}(s) = \left(1+\beta s\right)^{-\alpha} \tag{N1-6}$$

where shape parameter $\alpha$ and scale parameter $\beta$ related to the mean $\langle T\rangle$ and variance $\langle \delta T^2\rangle$ of $G(T)$ by

$$\alpha\beta = \langle T\rangle \text{ and } \alpha\beta^2 = \langle \delta T^2\rangle \tag{N1-7}$$

The series expansion of $\ln \hat{G}(s) = -\alpha \ln\left(1+\beta s\right)$ around $s=0$ is given by

$$\ln \hat{G}(s) = -\alpha\beta s + \frac{\alpha\beta^2}{2}s^2 + \cdots \qquad \left(s << \beta^{-1}\right) \tag{N1-8}$$

Using equation (N1-6), we can rearrange equation (N1-7) as

$$\ln \hat{G}(s) = -\langle T\rangle s + \frac{\langle \delta T^2\rangle}{2}s^2 + \cdots \qquad \left(s << \beta^{-1}\right) \tag{N1-9}$$

The equivalence between the right-hand-sides of equations (N1-4) and (N1-9) show that $\Phi(T)$ can be approximated by gamma distribution $G(T)$ at times long than $\beta = \langle \delta T^2 \rangle / \langle T \rangle$

When reaction times $\{\tau_1, \tau_2, \cdots, \tau_n\}$ are similarly distributed independent random variables, the mean total reaction time increases with the number of reaction steps, i.e., $\langle T \rangle = n\langle t \rangle$ where $\langle t \rangle \equiv n^{-1}\sum_{i=1}^{n}\langle t_i \rangle$. On the other hand, the randomness of total reaction time $T$ decreases with the number of elementary reactions composing the total reaction. The randomness of reaction time is often estimated by its relative fluctuation, which is given by

$$\frac{\langle \delta T^2 \rangle}{\langle T \rangle^2} = \frac{1}{n}\frac{\langle \delta t^2 \rangle}{\langle t \rangle^2} = \frac{1}{\langle T \rangle}\frac{\langle \delta t^2 \rangle}{\langle t \rangle} \tag{N1-10}$$

where $\langle \delta t^2 \rangle$ are defined by $\langle \delta t^2 \rangle \equiv n^{-1}\sum_{i=1}^{n}\langle \delta t_i^2 \rangle$. The relative variance of the total reaction time $T$ decreases with $n$ or $\langle T \rangle$, and becomes less than unity when the number $n$ of elementary steps is large enough as long as the mean $\langle t \rangle$ and variance $\langle \delta t^2 \rangle$ of individual reaction time exists. In the large $n$ limit, the gamma distribution approximation of $\Phi(T)$ approaches Gaussian with mean $\langle T \rangle$ and relative variance given in equation (N1-10) in accordance with Gaussian central limit theorem.

However, when the reaction times are strongly correlated, the relative variance $\langle \delta^2 T \rangle / \langle T \rangle^2$ is weakly dependent on $n$ or $\langle T \rangle$ (see Supplementary Note 6).

**Supplementary Note 2. Gamma Central Limit Theorem for Reaction Time Distribution of Arbitrary Multi-Channel Processes**

In this note, we will show that the reaction time distribution of a cell-state dependent rate process can be approximated by a super-Poisson gamma distribution. Let us consider a one-step reaction process with the rate coefficient $k(r)$ dependent on slow state variable $r$ of cell. For cells at state $r$, the reaction time distribution is given by $\varphi_r(T) = k(r)\exp[-k(r)T]$. For a given distribution $p(r)$ of cell state, the cell-state averaged reaction time distribution $\varphi(t)$ is given by $\varphi(T) = \int dr\varphi_r(T)p(r)$. Then $\varphi(T)$ can be approximated by a super-Poisson gamma distribution.

The proof of this statement is as follows. The Laplace transform of $\varphi(T)$ is given by

$$\hat{\varphi}(s) = \int \frac{1}{1+\tau(r)s} p(r)dr \tag{N2-1}$$

where $\tau(r)$ is defined by $\tau(r) \equiv k(r)^{-1}$. By expanding equation (N5-1) around $s=0$, we obtain

$$\hat{\varphi}(s) \cong 1 - \langle\tau\rangle s + \langle\tau^2\rangle s^2 + \cdots \tag{N2-2}$$

where $\langle\tau^n\rangle$ is defined by $\langle\tau^n\rangle = \int dr[k(r)]^{-n} p(r)$. The reaction time distribution given in equation (N2-2) approaches a super-Poisson gamma distribution, or a gamma distribution with the variance greater than square the mean. The Laplace transform of a gamma distribution with shape-scale parameters $(\alpha, \beta)$ can be expanded around $s=0$ as follows:

$$(1+s\beta)^{-\alpha} \cong 1 - \alpha\beta s + \frac{1}{2}\alpha(1+\alpha)\beta^2 s^2 + \cdots \qquad (s << \beta^{-1}) \tag{N2-3}$$

The equivalence between the right-hand-sides of equations (N2-2) and (N2-3) show that $\varphi(t)$ can be approximated by gamma distribution at times long than $\beta = \langle \delta t^2 \rangle / \langle t \rangle$. By comparing the series expansion in (N2-2) and (N2-3), we find the following relations: $\alpha\beta = \langle \tau \rangle$ and $\frac{1}{2}\alpha(1+\alpha)\beta^2 = \langle \tau^2 \rangle$. From these relations and equation (N1-7), we identify the mean reaction time $\langle T \rangle$ and the relative variance $\langle \delta T^2 \rangle / \langle T \rangle^2$ of reaction time as

$$\langle T \rangle = \langle \tau \rangle \qquad \text{(N2-4a)}$$

$$\frac{\langle \delta T^2 \rangle}{\langle T \rangle^2} = 1 + 2\eta_\tau^2 \qquad \text{(N2-4b)}$$

where $\eta_\tau^2$ denotes the relative variance of $\tau(r)$, i.e., $\eta_\tau^2 = \left( \langle \tau^2 \rangle - \langle \tau \rangle^2 \right) / \langle \tau \rangle^2$. As $\eta_\tau^2 > 0$, the relative variance given in equation (N2-4b) is always greater than unity. This result indicates that the reaction time distribution of a cell-state dependent one step process is a super-Poisson distribution.

**Supplementary Note 3. Model and Nomenclature**

In the present work, we define the signal transduction time, $t_{st}$, as the time delay between the onset of an environmental change and the first ensuing burst of target gene expression. $t_{st}$ is given by the sum of the target gene activation time, $t_{ga}$, or the time between the onset of the environmental switch and the target gene activation, and the first gene expression burst time, $\tau_b^{(1)}$, of the activated gene (see Fig. 1a). Then the distribution $\varphi_{st}(t)$ of the signal transduction time $t_{st}$ is given by the convolution of the distribution $\varphi_{ga}(t)$ of gene activation time $t_{ga}$ and the distribution $\varphi_b^{(1)}(t)$ of the first gene expression burst time, $\tau_b^{(1)}$. The gene activation and the first gene expression burst are multi-step, cell-state dependent non-Poisson processes whose RTDs can be modeled as gamma distributions (Supplementary Note 1). Their dynamics is far different from a one-step reaction process with a constant rate coefficient, for which the RTD is given by an exponential function.

The time delay between the successive target gene expression bursts[2], subsequent to the first gene expression burst, has a distinct probability distribution $\varphi_b(t)$, which usually has a far smaller mean value compared to $\varphi_{st}(t)$. In this case, the second or the higher moment of $\varphi_b(t)$ does not significantly affect the gene expression statistics so that $\varphi_b(t)$ can be modeled as a simple exponential function. On the other hand, when the target gene expression bursts occur on a time scale comparable to the signal transduction process, we model $\varphi_b(t)$ as a gamma distribution. In our model, the number of nascent proteins created per gene expression burst can be an arbitrary stochastic variable with finite mean and variance. However, according to refs.[3,4], it is assumed to follow a negative binomial distribution in our analysis of *E. coli* gene expression bursts in response to external signal.

The nascent proteins undergo maturation or post-translational modifications to become functional proteins. To describe the dynamics of the maturation process, we introduce the protein maturation time distribution $\varphi_m(t)$, or the distribution of time, $\tau_m$, taken for a nascent protein to become a mature protein. A representative example is the maturation of nascent fluorescent proteins. This maturation process occurs through the several post-translational modification steps: initial folding of nascent fluorescent protein to form a beta-barrel structure, internal cyclization to form a five-membered ring structure from the protein backbone, dehydration reaction that removes a water molecule from the ring structure, and the final oxidative reaction that converts the cyclized ring into the fully conjugated form[5]. The maturation time distribution varies depending on the type of fluorescent protein, which can be directly extracted from experimental data in literature[6] (Fig. 2a).

Finally, in our model, the lifetime of mature proteins can be an arbitrary stochastic variable whose probability distribution, $\varphi_d(t)$, depends on the protein degradation mechanism[1]. In the absence of such information, $\varphi_d(t)$ is often approximated by a simple exponential function, under the assumption that the protein degradation process has a single rate-determining elementary reaction step.

**Supplementary Note 4. Relationship between the rate-rate autocorrelation of immature protein burst and the signal transduction time distribution and gene expression burst interval distribution**

Let us consider a time series, $\{t_1^c, t_2^c, \cdots\}$, of reaction events, where $t_i^c$ denotes the time at which the $i$-th immature protein burst event is completed. The number $n(t)$ of immature protein burst events occurring in time interval $(0,t)$ is given by $\sum_{i=1}^{\infty}\Theta(t-t_i^c)$ where $\Theta(x)$ denotes the Heaviside step function. As the chemical reaction process is a stochastic process, the reaction times $\{t_i^c\}$ and $n(t)$ are random variables. The rate $R$ of immature protein burst events is defined as $R = dn(t)/dt$ . Noting that $d\Theta(t-t_i^c)/dt = \delta(t-t_i^c)$, where $\delta(x)$ is the Dirac delta function, we obtain

$$R(t) = \sum_{i=1}^{\infty}\delta(t-t_i^c) \tag{N4-1a}$$

By taking average of (N1-1) over the distribution of $\{t_i^c\}$, we obtain

$$\langle R(t)\rangle = \sum_{i=1}^{\infty}\langle\delta(t-t_i^c)\rangle = \sum_{i=1}^{\infty}\psi_i(t), \tag{N4-1b}$$

where $\psi_i(\tau_1)\left(=\langle\delta(\tau_1-t_i^c)\rangle\right)$ denotes the probability density of the time $t_i^c$ at which the $i$-th immature protein burst takes place. Since $t_i^c$ is the sum of the signal transduction time $t_{st}$, which is the sum of gene activation time and the first immature protein burst time, and the time taken for $i-1$ subsequent immature protein burst, the Laplace transform of $\psi_i(t)$ is given by $\hat{\psi}_i(s) = \hat{\varphi}_{st}(s)\hat{\varphi}_b(s)^{i-1}$. By substituting this result into the Laplace transform of equation (N4-1a)

$$\left\langle \hat{R}(s) \right\rangle = \sum_{i=1}^{\infty} \hat{\varphi}_{st}(s)\hat{\varphi}_b^{i-1}(s) = \frac{\hat{\varphi}_{st}(s)}{1-\hat{\varphi}_b(s)} \tag{N4-2}$$

Equation N4-2 follows because the signal transduction time distribution $\varphi_{st}(t)$, or distribution of time between the cell signal sensing and the first burst, differs from the distribution $\varphi_b(t)$ of time between successive bursts of immature proteins.

Let us now derive the corresponding expression of $\left\langle R(\tau_1)R(\tau_2) \right\rangle$, the TCF of the fluctuation in the rate of the immature protein burst events defined by

$$\left\langle R(\tau_1)R(\tau_2) \right\rangle = \left\langle \sum_{i=1}^{\infty} \sum_{\substack{k=1 \\ i \neq k}}^{\infty} \delta(\tau_1 - t_i^c)\delta(\tau_2 - t_k^c) \right\rangle \tag{N4-3}$$

Note here that $t_k^c$ is the time at which the *k*-th immature protein burst event is completed, so $t_k^c$ increases with *k*. Therefore, we obtain $\left\langle \delta(\tau_1 - t_i^c)\delta(\tau_2 - t_k^c) \right\rangle = 0$ for all *k* less than *i* when $\tau_2 > \tau_1$, we obtain the following equation from equation (N4-3)

$$\left\langle R(\tau_1)R(\tau_2) \right\rangle = \left\langle \sum_{i=1}^{\infty} \sum_{k=i+1}^{\infty} \delta(\tau_1 - t_i^c)\delta(\tau_2 - t_k^c) \right\rangle \quad (\tau_2 > \tau_1) \tag{N4-4}$$

This equation simply means that, if the *i*-th burst events occurs at $\tau_1$ and the *k*-th burst event occurs at a later time $\tau_2$, *k* should be greater than *i* for any time series of reaction events. Using the following property of the Dirac delta function, $f(\tau_1)\delta(\tau_1 - t_i) = f(t_i)\delta(\tau_1 - t_i)$, we rewrite equation (N4-4) as

$$\begin{aligned}\langle R(\tau_1)R(\tau_2)\rangle &= \left\langle \sum_{i=1}^{\infty}\sum_{k=i+1}^{\infty}\delta(\tau_1 - t_i^c)\delta(\tau_2-\tau_1-(t_k^c-t_i^c))\right\rangle \\ &= \left\langle \sum_{i=1}^{\infty}\sum_{m=1}^{\infty}\delta(\tau_1 - t_i^c)\delta(\tau_2-\tau_1-(t_{i+m}^c-t_i^c))\right\rangle\end{aligned} \tag{N4-5}$$

For our model, the magnitude of the time interval between the $i$-th burst event and the ($i$+$m$)-th burst event, $t_{i+m}^c - t_i^c$, is independent of the time $t_i^c$ at which the $i$-th burst event occurs.

Therefore, we can rewrite equation (N4-5) as

$$\begin{aligned}\langle R(\tau_1)R(\tau_2)\rangle &= \left\langle \sum_{i=1}^{\infty}\sum_{m=1}^{\infty}\delta(\tau_1 - t_i^c)\delta(\tau_2-\tau_1-(t_{i+m}^c-t_i^c))\right\rangle \\ &= \sum_{i=1}^{\infty}\left\langle \delta(\tau_1 - t_i^c)\right\rangle \sum_{m=1}^{\infty}\left\langle\delta(\tau_2-\tau_1-(t_{i+m}^c-t_i^c))\right\rangle\end{aligned} \tag{N4-6}$$

In equation (N4-6), $\left\langle\delta(\tau_2-\tau_1-(t_{i+m}^c-t_i^c))\right\rangle$ denotes the probability density $\psi_{m,i}(\tau_2-\tau_1)$ of the time, $\tau_2-\tau_1 = t_{i+m}^c - t_i^c$, between the $i$-th burst and ($i$+$m$)-th burst. Since $t_{i+m}^c - t_i^c$ is the sum of time required for $m$ subsequent immature protein burst, the Laplace transform of $\psi_{m,i}(t)$ is given by $\hat{\psi}_{m,i}(s) = \hat{\varphi}_b(s)^m$ for all $i$. Likewise $\psi_i(t)$, $\psi_{m,i}(t)$ satisfies the normalization condition, i.e., $\int_0^\infty dt\psi_{m,i}(t) = 1$, for any $i$ and $m$. In terms of $\psi_i(t)$ and $\psi_{m,i}(t)$, equation (N4-6) can be rewritten as

$$\langle R(\tau_1)R(\tau_2)\rangle = \sum_{i=1}^{\infty}\psi_i(\tau_1)\sum_{m=1}^{\infty}\psi_{m,i}(\tau_2-\tau_1) \tag{N4-7}$$

We can rewrite equation (N4-7) as

$$\langle R(\tau_1)R(\tau_2)\rangle = \langle R(\tau_1)\rangle\langle R'(\tau_2-\tau_1)\rangle \tag{N4-8}$$

where $\langle R(\tau)\rangle$ and $\langle R'(\tau)\rangle$ are defined by equation (N4-1b) and $\langle R'(\tau)\rangle = \sum_{m=1}^{\infty}\psi_{m,i}(\tau)$, respectively. $\langle R(\tau)\rangle$ is simply related to $\varphi_{st}(t)$ and $\varphi_b(t)$ in the Laplace domain as given in equation (N4-2). Since $\hat{\psi}_{m,i}(s) = \hat{\varphi}_b(s)^m$, $\langle R'(\tau)\rangle$ is related to $\varphi_b(t)$ only by

$$\left\langle \hat{R}'(s)\right\rangle \equiv \sum_{m=1}^{\infty}\hat{\psi}_{m,i}(s) = \frac{\hat{\varphi}_b(s)}{1-\hat{\varphi}_b(s)}. \tag{N4-9}$$

Under the stationary initial condition where our observation begins at a time evenly distributed between immature protein burst events and where immature protein burst is a stationary process, the mean rate $\langle R(t)\rangle_{st}$ of immature protein burst events becomes constant in time. $\langle R(t)\rangle_{st}$ is given by

$$\langle R(t)\rangle_{st} = \langle R\rangle = \langle \tau_b\rangle^{-1} \tag{N4-10}$$

where $\langle \tau_b\rangle$ denotes the mean burst interval defined by $\langle \tau_b\rangle = \int_0^{\infty} d\tau \varphi_b(\tau)\tau$ . This result indicates that the mean burst rate at long-time steady state is independent of the signal transduction time distribution. Equation (N4-10) can be obtained from the generalization of equation (N4-2) for the stationary initial condition, i.e.,

$$\left\langle \hat{R}(s)\right\rangle_{st} = \sum_{i=1}^{\infty}\hat{\varphi}_b^{st}(s)\hat{\varphi}_b^{i-1}(s) = \frac{\hat{\varphi}_b^{st}(s)}{1-\hat{\varphi}_b(s)} \tag{N4-11}$$

where $\hat{\varphi}_b^{st}(s)$ denotes the Laplace transform of the distribution $\varphi_b^{st}(t)$ of the first immature protein burst time under the stationary initial condition. It is known that $\varphi_b^{st}(t)$ is given by[7,8]

$$\varphi_b^{st}(t) = \langle \tau_b\rangle^{-1}\int_t^{\infty}\varphi_b(\tau)d\tau = \langle \tau_b\rangle^{-1}\left[1-\int_0^t \varphi_b(\tau)d\tau\right] \tag{N4-12}$$

whose Laplace transform is given by $\hat{\varphi}_b^{st}(s) = \left[1 - \hat{\varphi}_b(s)\right] / \left(s\langle\tau_b\rangle\right)$. Substituting the latter result into equation (N4-11), we obtain $\left\langle \hat{R}(s) \right\rangle_{st} = (s\left\langle \tau_b \right\rangle)^{-1}$ whose time domain expression yields equation (N4-10). Likewise, under the stationary initial condition, the autocorrelation function, $\left\langle R(\tau_1)R(\tau_2) \right\rangle$, of the rate of immature protein burst events is given by

$$\left\langle R(\tau_1)R(\tau_2) \right\rangle = \left\langle R'(\tau_2 - \tau_1) \right\rangle \left\langle R \right\rangle \tag{N4-13}$$

which can be obtained by replacing $\left\langle R(t) \right\rangle$ with $\left\langle R(t) \right\rangle_{st}$ or $\left\langle R \right\rangle \left( = \left\langle \tau_b \right\rangle^{-1} \right)$ in equation (N4-8).

**Supplementary Note 5. Steady-state Mean and Variance of Mature proteins**

For our model shown in Fig. 1a, the long-time limit of the mean protein number (equation 1a) is given by:

$$\langle n(\infty)\rangle = \lim_{t\to\infty}\int_0^t \langle R_T(\tau)\rangle P_m(t-\tau)d\tau \tag{N5-1}$$

where $\langle R_T(t)\rangle\left[=\langle b\rangle\langle R(t)\rangle\right]$ denotes the mean rate of immature protein production at time $t$. Here, $\langle b\rangle$ and $\langle R(t)\rangle$ represent the mean number of nascent proteins produced per burst and the mean rate of immature protein burst events at time $t$, respectively, given that the onset of environmental change occurs at time 0. $P_m(t)$ denotes the probability that an immature protein generates at time 0 has completed the maturation process before time $t$ and has survived as a mature protein as of time $t$. This probability is given by the convolution integral of the maturation time distribution, $\varphi_m(t)$, of protein and the survival probability, $S_d(t)$, of the mature protein (equation(2)). By applying the Tauberian theorem, the analytic expression of $\langle n(\infty)\rangle$ can be obtained, i.e.,

$$\langle n(\infty)\rangle = \lim_{s\to 0} s\langle b\rangle\langle \hat{R}(s)\rangle\hat{\varphi}_m(s)\hat{S}_d(s). \tag{N5-2}$$

In the above equation, $\lim_{s\to 0} s\langle \hat{R}(s)\rangle$ is the same as the long-time limit value $\left\langle R(\infty)\right\rangle$ of the mean rate of the protein burst events. Using equation (N4-2), we obtain the expression of $\lim_{s\to 0} s\langle \hat{R}(s)\rangle$ as follows:

$$\lim_{t\to\infty}\left\langle R(t)\right\rangle = \lim_{s\to 0} s\langle \hat{R}(s)\rangle = \lim_{s\to 0}\frac{s\hat{\varphi}_{st}\left(s\right)}{1-\hat{\varphi}_b\left(s\right)} = \frac{1}{\langle \tau_b\rangle} \tag{N5-3}$$

where $\langle \tau_b \rangle$ denotes the mean burst time interval. Equation (N5-3) follows because $\lim_{s\to 0}[1-\hat{\varphi}_b(s)]/s = \lim_{s\to 0}[\hat{\varphi}_b(0)-\hat{\varphi}_b(s)]/s = -\partial_s \hat{\varphi}_b(s)\big]_{s=0} = \int_0^\infty d\tau \varphi_b(\tau)\tau \left(\equiv \langle \tau_b \rangle\right)$ . Similarly, the small $s$ limit value of $\hat{S}_d(s)\left(=[1-\hat{\varphi}_b(s)]/s\right)$ is given by $\hat{S}_d(0) = \int_0^\infty dt S_d(t) = \langle \tau_d \rangle$ with $\langle \tau_d \rangle$ denoting the mean lifetime of the mature proteins. Noting also that the small $s$ limits of $\hat{\varphi}_m(s)$ yields $\hat{\varphi}_m(0) = \int_0^\infty dt \varphi_m(t) = 1$ due to the normalization condition, we obtain the expression for the steady-state mean number of mature proteins from equation (N5-2) as follows:

$$\langle n(\infty) \rangle = \frac{\langle b \rangle \langle \tau_d \rangle}{\langle \tau_b \rangle} \tag{N5-4}$$

Let us now obtain the expression of the steady-state variance in the mature protein number by considering the long-time limit of equation (1b) in the main manuscript. In the long-time steady-state, the TCF $\langle \delta R_T(\tau_2) \delta R_T(\tau_1) \rangle$ of the immature protein creation rate fluctuation given in equation (M1-22) becomes dependent only on the time difference, $\tau_2 - \tau_1$, i.e.,

$$\begin{aligned}
\langle \delta R_T(\tau_2) \delta R_T(\tau_1) \rangle &= \left(\langle b^2 \rangle - \langle b \rangle\right) \delta(\tau_2 - \tau_1) \langle R(\tau_1) \rangle + \langle b \rangle^2 \left[\langle R(\tau_2) R(\tau_1) \rangle - \langle R(\tau_2) \rangle \langle R(\tau_2) \rangle\right] \\
&= \left(\langle b^2 \rangle - \langle b \rangle\right) \delta(\tau_2 - \tau_1) \langle R \rangle + \langle b \rangle^2 \left[\langle R'(\tau_2 - \tau_1) \rangle \langle R \rangle - \langle R \rangle^2\right] \\
&\equiv C_{R_T}\left(\tau_2 - \tau_1\right)
\end{aligned} \tag{N5-5}$$

where $\langle R \rangle$ is the long-time limit of the mean protein burst rate, as defined in equation (N4-10) or (N5-3). Substituting $\langle \delta R_T(\tau_2) \delta R_T(\tau_1) \rangle = C_{R_T}\left(\tau_2 - \tau_1\right)$ into equation (1b) and taking the long-time limit of the resulting equation, we obtain

$$\sigma_n^2(\infty) = \langle n(\infty) \rangle + \int_0^\infty d\tau_2 \int_0^\infty d\tau_1 C_{R_T}\left(\tau_2 - \tau_1\right) P_m(\tau_2) P_m(\tau_1) \tag{N5-6}$$

By substituting the inverse Fourier transform representation of $C_{R_T}(\tau_2-\tau_1)=(2\pi)^{-1}\int_{-\infty}^{\infty}dwe^{iw(\tau_2-\tau_1)}\tilde{C}_{R_T}(w)$ into equation (N5-6), we obtain

$$\sigma_n^2(\infty)=\langle n(\infty)\rangle+\frac{1}{2\pi}\int_{-\infty}^{\infty}dw\tilde{C}_{R_T}\left(w\right)\left|\hat{P}_m(iw)\right|^2 \tag{N5-7}$$

where $\tilde{C}_{R_T}\left(w\right)$ denotes the Fourier transform of $C_{R_T}(t)$, defined by $\tilde{C}_{R_T}\left(w\right)=\int_{-\infty}^{\infty}dte^{-iwt}C_{R_T}(t)$. In equation (N5-7), $\hat{P}_m(s)$ denotes the Laplace transform of $P_m(t)$. From the definition of $P_m(t)$ given in equation (2) in main text, we obtain $\left|\hat{P}_m(iw)\right|^2=\left|\varphi_m(iw)\right|^2\left|S_d(iw)\right|^2$ using the convolution theorem. As $C_{R_T}(t)$ is an even function of time, $\tilde{C}_{R_T}\left(w\right)$ is related to the Laplace transform, $\hat{C}_{R_T}(s)=\int_0^{\infty}dte^{-st}C_{R_T}(t)$, of $C_{R_T}(t)$, by $\tilde{C}_{R_T}(w)=\hat{C}_{R_T}(iw)+\hat{C}_{R_T}(-iw)=2\,\mathrm{Re}\left[\hat{C}_{R_T}(iw)\right]$. Using these equations, we can rewrite equation (N5-7) as follows:

$$\sigma_n^2(\infty)=\langle n(\infty)\rangle+\frac{1}{\pi}\int_{-\infty}^{\infty}dw\,\mathrm{Re}\left[\hat{C}_{R_T}(iw)\right]\left|\hat{\varphi}_m(iw)\right|^2\left|\hat{S}_d(iw)\right|^2 \tag{N5-8}$$

The burst-interval and degradation-time distributions are often assumed to be exponential distributions, i.e., $\varphi_b\left(t\right)=ke^{-kt}$ and $\varphi_d\left(t\right)=\gamma e^{-\gamma t}$ with $k=1/\langle\tau_b\rangle$ and $\gamma=1/\langle\tau_d\rangle$. In this case, we have $\hat{\varphi}_b(iw)=k/(iw+k)$ and $\hat{\varphi}_d(iw)=\gamma/(iw+\gamma)$. However, the signal transduction time distribution $\varphi_{st}(t)$ and protein maturation time distribution $\varphi_m(t)$ deviate much from the simple exponential function.

Using equation (N5-4), we obtain the expression for the steady-state mean number of mature proteins is obtained as $\langle n(\infty)\rangle=\langle b\rangle k/\gamma$.

When the protein burst interval distribution follows the exponential function, $\varphi_b(t) = ke^{-kt}$, we have $\langle R \rangle = k$ and $\langle \delta R(\tau_2)\delta R(\tau_1) \rangle = 0$ so that $C_{R_T}(\tau)$ defined in (N5-5) reduces to $C_{R_T}(\tau) = \left(\langle b^2 \rangle - \langle b \rangle\right)\delta(\tau)k$ whose Fourier transform is given by $\tilde{C}_{R_T}(iw) = 2\,\mathrm{Re}\left[\hat{C}_{R_T}(iw)\right] = \left(\langle b^2 \rangle - \langle b \rangle\right)k$. In addition, when the protein lifetime distribution follow the exponential function $\varphi_d(t) = \gamma e^{-\gamma t}$, we have $S_d(t) = \int_t^\infty d\tau \varphi_d(\tau) = \exp(-\gamma t)$, whose Laplace transform is given by $\hat{S}_d(iw) = (iw + \gamma)^{-1}$. Substituting these results into equation (N5-8), we obtain the expression for the steady state variance of mature protein number as follows:

$$\sigma_n^{\,2}(\infty) = \langle n(\infty) \rangle + \left(\langle b^2 \rangle - \langle b \rangle\right)\frac{k}{2\pi}\int_{-\infty}^{\infty} \frac{\left|\hat{\varphi}_m(iw)\right|^2}{w^2 + \gamma^2} dw \qquad \text{(N5-9)}$$

In the present work, the protein maturation time distribution is modeled by a gamma distribution. In terms of the mean $\langle \tau_m \rangle$ and relative variance $\eta_m^2$ (i.e., $\eta_m^2 = \langle \delta\tau_m^{\;2} \rangle / \langle \tau_m \rangle^2$) of the maturation time, $\hat{\varphi}_m(iw)$ is given by , $\left(1 + iw\langle \tau_m \rangle \eta_m^2\right)^{-\frac{1}{\eta_m^2}}$ .Substituting this result into equation (N5-9), we obtain

$$\sigma_n^{\,2}(\infty) = \langle n(\infty) \rangle + \left(\langle b^2 \rangle - \langle b \rangle\right)\frac{k}{2\pi}\int_{-\infty}^{\infty} \frac{\left(1 + \left(\langle \tau_m \rangle \eta_m^2\right)^2 w^2\right)^{-\frac{1}{\eta_m^2}}}{\left(w^2 + \gamma^2\right)} dw \qquad \text{(N5-10)}$$

The integral in equation (N5-10) yields the following analytic expression:

$$\int_{-\infty}^{\infty} dw \frac{\left(1+\left(\langle\tau_m\rangle\eta_m^2\right)^2 w^2\right)^{-\frac{1}{\eta_m^2}}}{w^2+\gamma^2} = \pi\left(\frac{1}{\gamma}\left(\left(\langle\tau_m\rangle\eta_m^2\gamma\right)^2-1\right)^{-\frac{1}{\eta_m^2}} - \frac{\sqrt{\pi}\,{}_2F_1\left(\frac{1}{2},1;\frac{3}{2}-\frac{1}{\eta_m^2};\frac{1}{\left(\langle\tau_m\rangle\eta_m^2\gamma\right)^2}\right)}{\gamma^2\langle\tau_m\rangle\eta_m^2\Gamma\left(\frac{1}{\eta_m^2}\right)\Gamma\left(\frac{3}{2}-\frac{1}{\eta_m^2}\right)}\right)\mathrm{Sec}\left(\frac{\pi}{\eta_m^2}\right)$$

(N5-11)

In the small fluctuation limit, the maturation time distribution approaches a Dirac delta function, $\varphi_m(t)\to\delta\left(t-\langle\tau_m\rangle\right)$ (equivalently, $\hat{\varphi}_m(iw)=\left(1+\left(\langle\tau_m\rangle\eta_m^2\right)^2 w^2\right)^{-\frac{1}{\eta_m^2}}\to 1$ ) and the integral in equation (N5-10) yields $\int_{-\infty}^{\infty}\frac{1}{w^2+r^2}dw=\frac{\pi}{r}$. Substituting this result into equation (N5-10), we obtain the steady-state variance of the mature protein number in the small fluctuation limit.

$$\sigma_n^{\,2}(\infty)=\left\langle n(\infty)\right\rangle+\left(\left\langle b^2\right\rangle-\left\langle b\right\rangle\right)\frac{k}{2r} \qquad \text{(N5-12)}$$

In the opposite large fluctuation limit, $\eta_m^2\to\infty$ as well, we have $\hat{\varphi}_m(iw)=\left(1+\left(\langle\tau_m\rangle\eta_m^2\right)^2 w^2\right)^{-\frac{1}{\eta_m^2}}\to 1$ and recover (N5-12).

Finally, let us consider a more general model in which the mean burst rate $k$ of nascent protein and the decay rate $\gamma$ of the mature protein exhibit a significant cell-to-cell variation and follow arbitrary distributions across cells. For fixed values of $k$ and $\gamma$, the steady-state second moment $\left\langle n^2(\infty)\right\rangle_{k,\gamma}$ of the mature protein number is given by

$$\left\langle n^2(\infty)\right\rangle_{k,\gamma}=\left\langle n(\infty)\right\rangle_{k,\gamma}^2+\left\langle n(\infty)\right\rangle_{k,\gamma}+\left(\left\langle b^2\right\rangle-\left\langle b\right\rangle\right)\frac{k}{2\gamma} \qquad \text{(N5-13)}$$

where the first moments of mature protein number $\langle n(\infty)\rangle_{k,\gamma}$ in a cell with burst interval rate $k$ and the mean protein decay rate $\gamma$ is given by $\langle n(\infty)\rangle = \langle b\rangle k/\gamma$. By taking average of equation (N5-13) over the joint-distribution $p(k,\gamma)$ of $k$ and $\gamma$, we obtain

$$\left\langle \langle n^2(\infty)\rangle_{k,\gamma} \right\rangle = \langle b\rangle^2 \left\langle (k/\gamma)^2 \right\rangle + \langle b\rangle \langle k/\gamma\rangle + 2^{-1}\left(\langle b^2\rangle - \langle b\rangle\right)\langle k/\gamma\rangle \tag{N5-14}$$

where $\left\langle (k/\gamma)^n \right\rangle$ denotes $\int_0^\infty dk \int_0^\infty d\gamma (k/\gamma)^n p(k,\gamma)$. Equation (N5-14) immediately yields the following expression of the steady state variance:

$$\left\langle \langle n^2(\infty)\rangle_{k,\gamma} \right\rangle - \left\langle \langle n(\infty)\rangle_{k,\gamma}^2 \right\rangle = \langle b\rangle^2 \left[ \left\langle (k/\gamma)^2 \right\rangle - \langle k/\gamma\rangle^2 \right] + \langle b\rangle \langle k/\gamma\rangle + 2^{-1}\left(\langle b^2\rangle - \langle b\rangle\right)\langle k/\gamma\rangle$$

(N5-15)

**Supplementary Note 6. Mean- Variance Relationship of Multi-Step Processes**

Many biochemical processes consist of a large number of sequential elementary processes. In this note, we obtain the relationship between the mean and variance of the multi-step reaction time distribution and how it depends on the correlation among the reaction times of elementary reactions composing the multi-step reaction. For multi-step processes, the total reaction time is given by the sum of reaction times of the constituent elementary processes.

$$T_N = \sum_{i=1}^{N} t_i \tag{N6-1}$$

where $N$ and $t_i$ denote the number of elementary processes constituting the multi-step process and the reaction time of the $i$-th elementary process, respectively. Let us denote the mean total reaction time as $\langle T_N \rangle = \sum_{i=1}^{N} \langle t_i \rangle$ where $\langle t_i \rangle$ denotes the mean reaction time of the $i$-th elementary process. Then, the fluctuation, $\delta T_N \left( \equiv T_N - \langle T_N \rangle \right)$, of the total reaction time around its mean value is given by

$$\delta T_N = \sum_{i=1}^{N} \delta t_i \tag{N6-2}$$

with $\delta t_i = t_i - \langle t_i \rangle$ and the variance of $T_N$ is given by

$$\langle \delta T_N{}^2 \rangle = \sum_{i=1}^{N} \langle \delta t_i{}^2 \rangle + \sum_{i=1}^{N} \sum_{\substack{j=1 \\ i \neq j}}^{N} \langle \delta t_i \delta t_j \rangle \tag{N6-3}$$

When the reaction times of elementary processes are statistically independent random variables, or when correlation between the reaction times are negligible, the variance of the total reaction time linearly increases with the mean total reaction time. In this case, the second term on the r.h.s. of equation (N6-3) can be neglected ($\langle \delta t_i \delta t_j \rangle = 0$ , $i \neq j$), and the total mean and variance of the reaction time are given by $\langle T_N\rangle = N\langle t\rangle_1$, $\langle \delta T_N{}^2\rangle = N\sigma_t^2$ where $\langle t\rangle_1$ and $\sigma_t^2$ are defined by $\langle t\rangle = N^{-1}\sum_{i=1}^{N}\langle t_i\rangle$ and $\sigma_t^2 = N^{-1}\sum_{i=1}^{N}\langle \delta t_i^2\rangle$, respectively. Given that $\langle t\rangle_1$ and $\sigma_t^2$ rapidly approach *N*-independent constants as $N$ increases, the variance of the total reaction time is linearly proportional to its mean value, $\langle \delta T_N^2\rangle = c\langle T_N\rangle$ with $c = \langle \delta t^2\rangle / \langle t\rangle$. Therefore, the relative variance of the total reaction time inversely scales with the mean reaction time, i.e.,

$$\frac{\langle \delta T_N{}^2\rangle}{\langle T_N\rangle^2} = \frac{c}{\langle T_N\rangle} \qquad \text{(N6-4)}$$

This is the case for the fluorescent protein maturation process, which comprises a series of mutually independent, unimolecular processes (see Fig. 2a). Note that the relative fluctuation given in equation (N6-4) decreases with the mean total reaction time.

On the other hand, when the reaction times of the elementary processes are strongly correlated, the variance of the total reaction time exhibits a quadratic dependence on the mean total reaction time. In this case, the second term on the r.h.s. of equation (N6-3) is no longer negligible and can be written as $N(N-1)C_t$ where $N(N-1)$ and $C_t$ designate, respectively, the number of summands in the second term and the correlation between the reaction times of the elementary processes, defined by $C_t \equiv [N(N-1)]^{-1}\sum_{i=1}^{N}\sum_{\substack{j=1 \\ i\neq j}}^{N}\langle \delta t_i \delta t_j\rangle$. Then the variance of the total reaction time can be rewritten as $\langle \delta T_N{}^2\rangle = N\sigma_t^2 + N(N-1)\overline{C}_t$. Given that $C_t$ rapidly approaches an *N*-independent constant as $N$ increases, the variance given in equation (N6-5) is a quadratic function of the mean total reaction time, i.e.,

$$\langle \delta T_N{}^2\rangle = a\langle T_N\rangle^2 + b\langle T_N\rangle \qquad \text{(N6-5)}$$

where $a$ and $b$ are defined by $a = C_t / \langle t \rangle_1^2$ and $b = \sigma_t^2 / \langle t \rangle_1 - C_t$, respectively. This result shows that, when reaction times of elementary processes constituting the total process are strongly correlated, the variance exhibits a quadratic dependence on the mean total reaction time, as observed in gene activation processes (see Fig. 3i).

Such strong correlation between the reaction times of elementary processes originates from coupling between the reaction times and the heterogeneous cell environment. Let us consider the gene activation process consisting of *N* elementary processes whose reaction time distributions depend on statically heterogeneous environmental sate variables, $\Gamma$. If the constituent elementary processes are independent from each other for a given cell environmental state $\Gamma$, the Laplace transform of the total reaction time distribution is given by

$$\hat{\Phi}_T(s \mid \Gamma) = \prod_{i=1}^{n} \hat{\varphi}_i(s \mid \Gamma) \tag{N6-6}$$

where $\hat{\varphi}_i(s \mid \Gamma)$ denotes the Laplace transform of the reaction time distribution of the *i*-th elementary process in a cell at environmental state $\Gamma$. Then, the first two moments of the total reaction time distribution are obtained as

$$\langle T_N \rangle_\Gamma \left( \equiv \int_0^\infty dt \Phi_T(t \mid \Gamma) t \right) = \partial_s \hat{\Phi}_T(s \mid \Gamma) \Big|_{s=0} = \sum_{i=1}^{N} \langle t_i \rangle_\Gamma \tag{N6-7}$$

and

$$\langle T_N^2 \rangle_\Gamma \left( \equiv \int_0^\infty dt \Phi_T(t \mid \Gamma) t^2 \right) = \partial_s^2 \hat{\Phi}_T(s \mid \Gamma) \Big|_{s=0} = \sum_{i=1}^{N} \langle t_i^2 \rangle_\Gamma + \sum_{i=1}^{N} \sum_{\substack{j=1 \\ j \neq i}}^{N} \langle t_i \rangle_\Gamma \langle t_j \rangle_\Gamma \tag{N6-8}$$

where $\langle t_i \rangle_\Gamma$ and $\langle t_i^2 \rangle_\Gamma$ denote the first two moments of reaction time for the *i*-th elementary process in a cell under environmental state $\Gamma$. When the environmental state is fixed, the mean and variance of the total reaction time are given by $\langle T_N\rangle_\Gamma = N\langle t\rangle_\Gamma$ and $\langle \delta T_N^2\rangle_\Gamma = N\langle \delta t^2\rangle_\Gamma$ where $\langle t\rangle_\Gamma$ and $\langle \delta t^2\rangle_\Gamma$ are defined by $\langle t\rangle_\Gamma = N^{-1}\sum_{i=1}^{N}\langle t_i\rangle_\Gamma$ and $\langle \delta t^2\rangle_\Gamma = N^{-1}\sum_{i=1}^{N}\langle t_i^2\rangle_\Gamma - \langle t_i\rangle_\Gamma^2$, respectively. Given that $\langle t\rangle_\Gamma$ and $\langle \delta t^2\rangle_\Gamma$ rapidly approach $N$-independent constants as $N$ increases, the variance of the total reaction time is linearly proportional to its mean value, $\langle \delta T_N^2\rangle_\Gamma = c_\Gamma \langle T_N\rangle_\Gamma$ with $c_\Gamma = \langle \delta t^2\rangle_\Gamma / \langle t\rangle_\Gamma$.

On the other hand, the variance of the total reaction time exhibits a quadratic dependence on the mean total reaction time when the environmental state is distributed. In this case, the environmental state average of equations (N6-7) and (N6-8) are given by $\langle\langle T_N\rangle_\Gamma\rangle = \sum_{i=1}^{N}\langle\langle t_i\rangle_\Gamma\rangle$ and $\langle\langle T_N^2\rangle_\Gamma\rangle = \sum_{i=1}^{N}\langle\langle t_i^2\rangle_\Gamma\rangle + \sum_{i\neq j}\langle\langle t_i\rangle_\Gamma\langle t_j\rangle_\Gamma\rangle$, respectively. Then the variance of the total reaction time can be written as

$$\langle\langle \delta T_N^2\rangle_\Gamma\rangle = N\left(\langle\langle t^2\rangle_\Gamma\rangle - \langle\langle t\rangle_\Gamma\rangle^2\right) + N(N-1)\left(\langle\langle t\rangle_\Gamma\langle t'\rangle_\Gamma\rangle - \langle\langle t\rangle_\Gamma\rangle\langle\langle t'\rangle_\Gamma\rangle\right) \tag{N6-9}$$

where $\langle\langle t^n\rangle_\Gamma\rangle$ and $\langle\langle t\rangle_\Gamma\langle t'\rangle_\Gamma\rangle - \langle\langle t\rangle_\Gamma\rangle\langle\langle t'\rangle_\Gamma\rangle$ are defined by $\langle\langle t^n\rangle_\Gamma\rangle \equiv N^{-1}\sum_{i=1}^{N}\langle\langle t_i^n\rangle_\Gamma\rangle$ and $\langle\langle t\rangle_\Gamma\langle t'\rangle_\Gamma\rangle - \langle\langle t\rangle_\Gamma\rangle\langle\langle t'\rangle_\Gamma\rangle \equiv [N(N-1)]^{-1}\sum_{i\neq j}\left(\langle\langle t_i\rangle_\Gamma\langle t_j\rangle_\Gamma\rangle - \langle\langle t_i\rangle_\Gamma\rangle\langle\langle t_j\rangle_\Gamma\rangle\right)$, respectively. Given that both quantities rapidly approach $N$-independent constant as $N$ increases, the variance of the total reaction time becomes a quadratic function of its mean, which is evident from the following equation obtained by substituting $N = \langle\langle T_N\rangle_\Gamma\rangle / \langle\langle t\rangle_\Gamma\rangle$ into equation (N6-9):

$$\left\langle\langle\delta T_N^2\rangle_\Gamma\right\rangle=\left\langle\langle T_N\rangle_\Gamma\right\rangle\frac{\left\langle\langle t^2\rangle_\Gamma\right\rangle-\left\langle\langle t\rangle_\Gamma\right\rangle^2-\left\langle\langle t\rangle_\Gamma\langle t'\rangle_\Gamma\right\rangle+\left\langle\langle t\rangle_\Gamma\right\rangle\left\langle\langle t'\rangle_\Gamma\right\rangle}{\left\langle\langle t\rangle_\Gamma\right\rangle}$$
$$+\left\langle\langle T_N\rangle_\Gamma\right\rangle^2\frac{\left\langle\langle t\rangle_\Gamma\langle t'\rangle_\Gamma\right\rangle-\left\langle\langle t\rangle_\Gamma\right\rangle\left\langle\langle t'\rangle_\Gamma\right\rangle}{\left\langle\langle t\rangle_\Gamma\right\rangle^2}. \tag{N6-10}$$

This result confirms the quadratic mean-variance relationship of the waiting time distribution under environmental fluctuations.

The quadratic mean-variance relationship of a waiting time distribution is a universal feature of dynamical systems with heterogeneous state parameters. For example, let us consider a particle undergoes diffusion under a constant force field in the one-dimensional space, whose probability distribution $p(x,t)$ at time $t$ satisfies the following Fokker–Planck equation:

$$\frac{\partial p}{\partial t}=D\frac{\partial^2 p}{\partial x^2}-v\frac{\partial p}{\partial x} \tag{N6-11}$$

where $D$ and $v$ denote the diffusion coefficient and the drift velocity induced by the constant force, respectively, and the system is subject to absorbing boundary at $x=L$.

It is well known that the first-passage time distribution, $f(t)$, of the particle initially located at $x=0$ to the absorbing boundary at $x=L$ follows an inverse-Gaussian distribution, $f_v(t)\sim IG\left(\frac{L}{v},\frac{L^2}{2D}\right)$. The mean and variance of the first passage time under a constant force field are given by $\langle t\rangle_v=\frac{L}{v}$, $\langle\delta t^2\rangle_v=\frac{2DL}{v^3}$. If the external force or drift velocity $v$ are statically distributed, the first passage time distribution $f(t)$ is given by $\langle f(t)\rangle=\int f_v(t)p(v)dv$ where $p(v)$ denotes the distribution of drift velocity, and the mean and variance are then given by

$$\langle\langle t\rangle_\nu\rangle = L\left\langle\frac{1}{\nu}\right\rangle, \quad \text{(N6-12a)}$$

$$\langle\langle \delta t^2\rangle_\nu\rangle = L^2\left(\left\langle\frac{1}{\nu^2}\right\rangle - \left\langle\frac{1}{\nu}\right\rangle^2\right) + 2DL\left\langle\frac{1}{\nu^3}\right\rangle. \quad \text{(N6-12b)}$$

Substituting $L = \langle\langle t\rangle_\nu\rangle / \langle\nu^{-1}\rangle$ into Eq. (N6-12b), we obtain

$$\langle\langle \delta t^2\rangle_\nu\rangle = \langle\langle t\rangle_\nu\rangle^2 \frac{\langle\nu^{-2}\rangle - \langle\nu^{-1}\rangle^2}{\langle\nu^{-1}\rangle^2} + 2D\langle\langle t\rangle_\nu\rangle \frac{\langle\nu^{-3}\rangle}{\langle\nu^{-1}\rangle}. \quad \text{(N6-13)}$$

This result shows that, even in a simple drift–diffusion system, static disorder in the external force can generate a quadratic dependence of the variance on the mean, analogous to the behavior observed in correlated multi-step gene-activation processs.

**Supplementary Note 7. Previous Theories and Models for Delayed Reaction Dynamics**

The chemical kinetics and the chemical master equation approaches[9] employed in the conventional systems biology provide ideal frameworks for description of chemical dynamics of reaction networks under homogeneous environments. However, these classical approaches cannot provide an accurate description for complex dynamics of intracellular reaction networks. This is not only because these reaction networks are complex and comprise a large number of elementary reactions, but also because the rate coefficients of each intracellular reaction processes are not constants but stochastic variables that differ from cell to cell and fluctuate over time, due to their coupling to dynamically heterogeneous cell environment[10-14].

Significant advances have been made in the quantitative description of the complex dynamics of reaction networks in living cells. Early this century, the delay chemical master equation (dCME) was proposed to describe the stochastic oscillation in gene regulation arising from delayed protein degradation processes[15]. This pioneering work, however, assumed that gene expression, composed of a series of enzyme reactions, is a simple Poisson process with a constant rate coefficient and it employed an unrealistic description of protein degradation with a constant time delay. Later, dCME was extended to incorporate a distribution of the time delay parameter[16] . However, even this extended dCME cannot accurately describe a birth-death dynamics of product molecules with a given lifetime distribution. As a matter of fact, an accurate chemical kinetics or chemical master equation has yet to be known for molecules with a general non-exponential lifetime distribution. Recently, the Chemical Fluctuation Theorem (CFT) was derived, which provides accurate analytic results for the time-dependent mean and variance of the number of product molecules with arbitrary lifetime distribution and product creation dynamics[1,17]. The CFT provides a successful quantitative explanation of the mean and variance of mRNA and protein levels across diverse gene expression systems[1,17]. The CFT provides a successful quantitative explanation of the steady-state mean and variance of mRNA and protein levels across diverse gene expression systems[1,17,18]. In principle, the CFT can also provide an accurate description for nonstationary, stochastic dynamics of reaction networks in living cells.

Despite these advances, however, a quantitative understanding of cell signal transduction and adaptive gene expression dynamics remains a formidable task. This difficulty arises because cell adaptation involves nonstationary stochastic processes whose dynamics markedly differ from the steady-state gene expression processes. In particular, cell adaptation invariably involves a strongly nonstationary process preceding target gene expression, which comprises environmental signal sensing, signal propagation, and target gene activation. Nascent proteins generated from gene expression undergo delayed maturation process, further compounding the complexity of adaptive cell dynamics.

Recently, an attempt was made to extend the CFT to describe time-delayed cell signaling dynamics[19]. However, this initial model relies on unrealistic assumptions, for example, that it merges signal-induced gene activation and mature protein production into a single process. Although this process is described by an arbitrary waiting time distribution, it assumes that after each round of mature protein production, the gene’s active state is reset, and the next round of gene activation occurs only for the subsequent protein synthesis. Together with signaling initiations following a stationary Poisson process, this assumption leads to the erroneous prediction that the mean and variance of the mature protein number are identical. In reality, once a signaling cascade is initiated, downstream processes such as transcription, translation, and protein maturation evolve continuously over time, rather than repeatedly resetting after each round[20,21]. To date, no existing model accounts for both signal transduction and protein maturation, nor has succeeded in providing a quantitative explanation of cell adaptation dynamics in response to environmental changes[22].

#### Supplementary Note 8. Stochastic simulation methods

Stochastic simulations of signal-induced fluorescent reporter expression were performed using a custom C program parallelized with MPI. Each single-cell trajectory consisted of a sequence of bursts of immature fluorescent protein creation occurring at $t_i^c$. The time, $t_1^c$, of the first burst time, measured from the onset of the external signal at time zero, was drawn from the distribution of the signal transduction time: either a gamma distribution (Fig. 1c-i and the lower panels of Fig. 2e-i) or a convolution of two gamma distributions (the upper panels of Fig. 2e-i). Successive burst intervals were sampled from an exponential distribution given by $\varphi_b(t) = \langle \tau_b \rangle^{-1} e^{-t/\langle \tau_b \rangle}$. At each burst time, $t_i^c$, the burst size, $b_i$, was sampled from a negative binomial distribution, $NB(5, 0.5)$. For each immature protein produced in a burst, the maturation time was sampled from the best-fit gamma distribution corresponding to the fluorescent protein used (Supplementary Fig. 2), while the protein lifetime was sampled independently from an exponential distribution given by $\varphi_d(t) = \langle \tau_d \rangle^{-1} e^{-t/\langle \tau_d \rangle}$.

To calculate the time-dependent number of mature proteins, $N_m(t)$, for the *m*th trajectory, the observation time window [0, *T*] was discretized into $N_{bin}$ bins of width $0.3\langle \tau_b \rangle$. At each time bin, $N_m(t)$ was calculated as the difference between the cumulative number of maturation events and the cumulative number of degradation events occurring since time zero. Using MPI_Reduce, values of $N_m(t)$ and $N_m^2(t)$ were aggregated across trajectories to obtain the ensemble-averaged quantities, including the mean and variance of the mature protein count. Parameter values used in the simulations are provided in Figs 1 and 2.

**Supplementary Note 9. Extraction of maturation time distributions for various fluorescent reporter proteins.**

The maturation time distribution, $\varphi_m(t)$, for a given fluorescent protein can be obtained from translation-stopped experiments[6]. Following chloramphenicol treatment of bacterial cells during gene expression, any subsequent increase in single-cell fluorescence intensity can be attributed to the maturation of proteins that were already synthesized prior to the treatment. These fluorescence intensity profiles were converted into the normalized fraction of immature proteins. However, the normalized fraction should not be interpreted as the survival probability, $S_m(t)\left[=\int_t^\infty d\tau\varphi_m(\tau)\right]$, of the maturation time because all immature proteins in cells are not simultaneously produced upon chloramphenicol treatment. Instead, we have to consider the following survival probability, $S_m^{st}(t)\left[=\int_t^\infty d\tau\varphi_m^{st}(\tau)\right]$, which is defined with the maturation time distribution, $\varphi_m^{st}(t)$, under stationary initial condition that the observation starts at an arbitrarily chosen time after maturation processes reach their stationary state:

$$S_m^{st}(t)=\int_t^\infty d\tau\varphi_m^{st}(\tau)=\frac{1}{\langle\tau_m\rangle}\int_t^\infty d\tau S_m(\tau) \qquad \text{(N9-1)}$$

with $\langle\tau_m\rangle$ denoting the mean maturation time.

Assuming that the maturation time distribution is given by $\varphi_m(t)=e^{-t/b_m}t^{a_m-1}\big/\Gamma(a_m){b_m}^{a_m}$ with $\langle\tau_m\rangle=a_m b_m$, $S_m^{st}(t)$ reads as

$$S_m^{st}(t)=\frac{1}{\langle\tau_m\rangle}\int_t^\infty d\tau\frac{\Gamma(a_m,\tau/b_m)}{\Gamma(a_m)}, \qquad \text{(N9-2).}$$

where $\Gamma(a)$ and $\Gamma(a,z)$ respectively denote the gamma function and the upper incomplete gamma function, defined by $\Gamma(a) = \int_0^\infty dt e^{-t} t^{a-1}$ and $\Gamma(a,z) = \int_z^\infty dt e^{-t} t^{a-1}$. The first-order and second-order moments, $\langle \tau_m^n \rangle^{st} \left[ = \int_0^\infty dt \varphi_m^{st}(t) t^n \right]$ ($n$ = 1,2), of $\varphi_m^{st}(t)$ can then be obtained by using equation (N9-2):

$$\begin{aligned}\langle \tau_m \rangle^{st} &= \int_0^\infty dt\, t \varphi_m^{st}(t) = -\int_0^\infty dt\, t \partial_t S_m^{st}(t) = -t S_m^{st}(t)\Big|_0^\infty + \int_0^\infty dt S_m^{st}(t) \\ &= \int_0^\infty dt S_m^{st}(t) = \frac{b_m(a_m+1)}{2},\end{aligned} \qquad \text{(N9-3a)}$$

$$\begin{aligned}\langle \tau_m^2 \rangle^{st} &= \int_0^\infty dt\, t^2 \varphi_m^{st}(t) = -\int_0^\infty dt\, t^2 \partial_t S_m^{st}(t) = -t^2 S_m^{st}(t)\Big|_0^\infty + 2\int_0^\infty dt\, t S_m^{st}(t) \\ &= 2\int_0^\infty dt\, t S_m^{st}(t) = \frac{b_m^2(a_m+2)(a_m+1)}{3},\end{aligned} \qquad \text{(N9-3b)}$$

where we have used the fact that $S_m^{st}(t=0) = 1$ and $S_m^{st}(t \to 0) \to 0$.

Finally, $a_m$ and $b_m$ can be determined using equations (N9-3a) and (N9-3b) instead of finding the best fit of equation (N9-2) to the immature-protein fraction data. Using the numerical representation, $S_{m,\exp}^{st}(t)$, interpolating the immature-protein fraction data for a given fluorescent protein, $\langle \tau_m \rangle_{\exp}^{st}$ and $\langle \tau_m^2 \rangle_{\exp}^{st}$ can be calculated as $\langle \tau_m \rangle_{\exp}^{st} = \int_0^\infty dt S_{m,\exp}^{st}(t)$ and $\langle \tau_m^2 \rangle_{\exp}^{st} = 2\int_0^\infty dt\, t S_{m,\exp}^{st}(t)$. Substituting the values of $\langle \tau_m \rangle_{\exp}^{st}$ and $\langle \tau_m^2 \rangle_{\exp}^{st}$ into equations (N9-3a) and (N9-3b), and solving the resulting equations, we can obtain the values of $a_m$ and $b_m$. The resulting profiles for $S_m^{st}(t)$ were found to be well superimposed on the immature-protein fraction data (Supplementary Fig. 2). In the present analysis, fluorescent proteins that did not reach a saturation regime within the experimental time window, codon-optimized variants, and second-valine fluorescent proteins were excluded.

**Supplementary Note 10. Data sources and preprocessing for quantitative gene expression analysis.**

The time-lapse microscopy data shown in Fig. 3b, c were obtained from a previous study[22]. In this experiment, colonies of *E. coli* were exposed to one of three antibiotic stresses: tetracycline (TET), trimethoprim (TMP), or nitrofurantoin (NIT). Under the TET stress condition, time-lapse YFP[23] expression for eight promoters (*dnaK, cspA, ydiU, ahpC, iscR, nrdH, rpsA,* and *rpmE*) and time-lapse CFP[24] expression for one promoter (*iscR*) were recorded. Under the TMP stress condition, time-lapse YFP expression for twelve promoters (*gadW, gadA, folA, recA, fpr, purT, purM, ldhA, guaB, gadB, osmC*, and *dps*) was recorded. Under the NIT stress condition, time-lapse YFP expression for four promoters (*fpr, ybjC, recA*, and *cysK*) was recorded. Each experiment was repeated up to four times.

We applied an inclusion criterion to ensure reliable estimation of the mean and variance of the mature protein expression level, retaining only datasets for which the number of cells at the final observation time exceeded 100. As a result, all NIT-stress datasets were excluded from further analysis. In addition, time-lapse YFP expression data for *recA* under the TMP stress condition were excluded due to excessive noise in the profiles. The remaining datasets include $dnaK_{1,2}$, $cspA_{2}$, $ydiU_{1,2}$, $iscR_{1,2}$, $nrdH_{1,2}$, $rpsA_{1,2}$, and $rpmE_{1,2}$, under the TET stress condition, and $gadA_{2}$, $folA_{1,2}$, $purM_{1,2}$, and $guaB_{1,2}$ under the TMP stress condition. Subscript labels attached to promoter names indicate one of two independent datasets collected from different microcolonies. To isolate stress-induced responses, the raw mean and variance of the fluorescence reporter intensity were baseline-shifted by subtracting an offset.

The IPTG-induced gene expression data shown in Fig. 3e were obtained from *E. coli* cells carrying plasmid-integrated sfGFP gene. Detailed experimental procedures are described in Experimental Methods. For each plasmid type (pAISA1, pAISA3, and pAISA5), which differ in their UTR sequences and thus exhibit different translational efficiencies for sfGFP expression, the mean and variance of the mature sfGFP copy number were averaged over three independent experimental repeats at each measurement time point. To isolate IPTG-induced responses, the raw mean and variance of the mature sfGFP copy number were baseline-shifted by subtracting an offset.

**Supplementary Note 11. Quantitative analyses of signal-induced gene expression data.**

To perform a quantitative analysis of the experimental data shown in Fig. 3b, c, we need explicit expressions for the time-dependent mean and variance, $\langle n(t)\rangle$ and $\sigma_n^2(t)\left[\equiv\langle\delta n^2(t)\rangle\right]$, of the mature-protein copy number, $n$. For our signal propagation-gene expression network model characterized by reaction time distributions, rather than rate constants, the Laplace-domain expressions are analytically tractable because time convolutions become products in this domain. Applying the Laplace transform to equation (1a) given in the main text, we obtain

$$\langle\hat{n}(s)\rangle=\langle b\rangle\langle\hat{R}(s)\rangle\hat{P}_m(s), \tag{N11-1}$$

where $\hat{f}(s)$ denotes the Laplace transform of $f(t)$, defined by $\hat{f}(s)=\mathcal{L}[f(t)]=\int_0^\infty dte^{-st}f(t)$. In equation (N11-1), $\langle b\rangle$, $\langle\hat{R}(s)\rangle$, and $\hat{P}_m(s)$ respectively denote the mean number of immature proteins produced per burst, the Laplace transform of the mean burst rate, $\langle R(t)\rangle$, and the Laplace transform of $P_m(t)$, defined by $P_m(t)=\int_0^t\varphi_m(\tau)S_d(t-\tau)d\tau$ with $S_d(t)$ being the survival probability of mature proteins [equation (2)].

In our quantitative analysis, the maturation-time distribution is modeled as a multi-exponential distribution,

$$\varphi_m(t)=\sum_{i=1}^{n}C_ie^{-a_it}, \tag{N11-2}$$

with $C_i$ and $a_i$ being constants. With the assumption that protein lifetimes follow an exponential distribution given by $\varphi_d(t)=\gamma e^{-t\gamma}$ (Supplementary Fig. 4), $\hat{P}_m(s)$ can be expressed as

$$\hat{P}_m(s) = \hat{S}_d(s)\hat{\varphi}_m(s) = \frac{1}{s+\gamma}\sum_{i=1}^{n}\frac{C_i}{s+a_i}, \tag{N11-3}$$

the inverse Laplace transformation of which yields

$$P_m(t) = \sum_{i=1}^{n}\frac{C_i}{a_i-\gamma}\left(e^{-\gamma t} - e^{-a_i t}\right). \tag{N11-4}$$

To go further, we need the Laplace-domain expression of the second-order moment, $\langle n^2(t)\rangle$, of the mature protein number [equation (M1-17)], which is reproduced for convenience below.

$$\begin{aligned}\left\langle n(t)^2\right\rangle &= \langle n(t)\rangle + \langle b(b-1)\rangle\int_0^t d\tau\left\langle R(\tau)\right\rangle P_m(t-\tau)^2 \\ &+ \langle b\rangle^2\int_0^t d\tau_2\int_0^t d\tau_1\left\langle R(\tau_1)R(\tau_2)\right\rangle P_m(t-\tau_1)P_m(t-\tau_2),\end{aligned} \tag{N11-5}$$

where $\langle b(b-1)\rangle$ can be expressed as $\langle b(b-1)\rangle = \langle b\rangle^2(1+1/m)$ with $m(>0)$ being the parameter characterizing the deviation from Poisson statistics; the large-$m$ limit corresponds to Poisson. On the r.h.s. of equation (N11-5), the second term can then be written in the Laplace domain as

$$\langle b\rangle^2(1+1/m)\left\langle\hat{R}(s)\right\rangle\mathcal{L}\left[P_m^2(t)\right]. \tag{N11-6}$$

In equation (N11-6), we need the expression of $P_m^2(t)$, which can be calculated using equation (N11-4) as

$$P_m^2(t) = \sum_{i=1}^{n}\sum_{k=1}^{n}\frac{C_i}{a_i-\gamma}\frac{C_k}{a_k-\gamma}\left(e^{-2\gamma t} - e^{-(a_i+\gamma)t} - e^{-(a_k+\gamma)t} + e^{-(a_i+a_k)t}\right), \tag{N11-7}$$

the Laplace transform of which is given by

$$\mathcal{L}\left[P_m^2(t)\right]=\sum_{i=1}^{n}\sum_{k=1}^{n}\frac{C_i}{a_i-\gamma}\frac{C_k}{a_k-\gamma}\left(\frac{1}{s+2\gamma}-\frac{1}{s+a_i+\gamma}-\frac{1}{s+a_k+\gamma}+\frac{1}{s+a_i+a_k}\right). \quad \text{(N11-8)}$$

In addition, on the r.h.s. of equation (N11-5), the third term can be rewritten as

$$\begin{aligned}&\langle b\rangle^2\int_0^t d\tau_2\int_0^t d\tau_1\left\langle R(\tau_1)R(\tau_2)\right\rangle P_m(t-\tau_1)P_m(t-\tau_2)\\&=2\langle b\rangle^2\int_0^t d\tau_1\int_{\tau_1}^t d\tau_2\left\langle R'(\tau_2-\tau_1)\right\rangle\left\langle R(\tau_1)\right\rangle P_m(t-\tau_1)P_m(t-\tau_2)\\&=2\langle b\rangle^2\int_0^t d\tau_1\left\langle R(\tau_1)\right\rangle P_m(t-\tau_1)\int_0^{t-\tau_1}dx\left\langle R'(x)\right\rangle P_m(t-\tau_1-x),\end{aligned} \quad \text{(N11-9)}$$

where the second equality follows from exchanging the order of integration and equation (N4-8) with $\langle R'(t)\rangle$ defined in equation (N4-9). From the last equality in equation (N11-9), $\langle b\rangle\int_0^t d\tau\left\langle R'(\tau)\right\rangle P_m(t-\tau)$ corresponds to the mean mature-protein number, $\langle n'(t)\rangle$, under synchronized initial condition that the distribution of the first protein-burst time is identical to the distribution, $\varphi_b(t)$, of the time interval between successive protein bursts. Based on this identification, equation (N11-9) can be expressed as

$$2\left\langle b\right\rangle\int_0^t d\tau_1\left\langle R(\tau_1)\right\rangle P_m(t-\tau_1)\left\langle n'(t-\tau_1)\right\rangle \quad \text{(N11-10)}$$

Substituting equation (N11-4) into equation (N11-10) and performing the Laplace transformation of the resulting equation, we obtain

$$2\left\langle b\right\rangle\left\langle\hat{R}(s)\right\rangle\left(\sum_{i=1}^{n}\frac{C_i}{a_i-\gamma}\left(\langle\hat{n}'(s+\gamma)\rangle-\langle\hat{n}'(s+a_i)\rangle\right)\right), \quad \text{(N11-11)}$$

where $\langle\hat{n}'(s)\rangle$ is given by $\langle b\rangle\langle\hat{R}'(s)\rangle\hat{P}_m(s)$.

For the quantitative analysis of the antibiotic stress-induced gene expression data (Fig. 3b, c), the distributions of the gene activation time and the ensuing first protein burst time are modelled as a gamma distribution: $\hat{\varphi}_{ga}(s)=[1+s\langle t_{ga}\rangle\eta_{ga}^2]^{-1/\eta_{ga}^2}$ and. $\hat{\varphi}_b^{(1)}(s)=[1+s\langle\tau_b^{(1)}\rangle\eta_{fb}^2]^{-1/\eta_{fb}^2}$ in the Laplace domain. The Laplace transform of the signal-transduction time distribution is then given by $\hat{\varphi}_{st}(s)=\hat{\varphi}_{ga}(s)\hat{\varphi}_b^{(1)}(s)$ $=[1+s\langle t_{ga}\rangle\eta_{ga}^2]^{-1/\eta_{ga}^2}[1+s\langle\tau_b^{(1)}\rangle\eta_{fb}^2]^{-1/\eta_{fb}^2}$ . Together with the burst-interval time distribution assumed to be an exponential distribution, given by $\varphi_b(t)=ke^{-kt}$ with $k=1/\langle\tau_b\rangle$, equations (N4-2) and (N4-9) read as

$$\langle\hat{R}(s)\rangle=\frac{\hat{\varphi}_{st}(s)}{1-\hat{\varphi}_b(s)}=\frac{[1+s\langle t_{ga}\rangle\eta_{ga}^2]^{-1/\eta_{ga}^2}[1+s\langle\tau_b^{(1)}\rangle\eta_{fb}^2]^{-1/\eta_{fb}^2}}{1-\dfrac{k}{s+k}}, \tag{N11-12a}$$

$$\langle\hat{R}'(s)\rangle=\frac{\hat{\varphi}_b(s)}{1-\hat{\varphi}_b(s)}=\frac{k}{s}. \tag{N11-12b}$$

For the antibiotic stress-response experiments at 30 °C, YFP (VenusNB) and CFP (mCerulean) were used as a fluorescent reporter. For numerical convenience, their maturation time distributions were assumed to be a biexponential function, i.e., equation (N11-2) with $n=2$, $C_i=a_ip_i$, and $p_1+p_2=1$, which well represents the time profiles for the normalized fraction of immature proteins for VenusNB and mCerulean (Supplementary Fig. 2). For this maturation-time distribution model, the analytic expression of $S_m^{st}(t)$ is given by [see equation (N9-1) in Supplementary Note 9]

$$\begin{aligned}S_m^{st}(t)&=\frac{1}{\langle\tau_m\rangle}\int_t^\infty d\tau S_m(\tau)=\frac{1}{\langle\tau_m\rangle}\int_t^\infty d\tau\left[p_1e^{-a_1\tau}+p_2e^{-a_2\tau}\right]\\&=\frac{p_1e^{-a_1t}/a_1+p_2e^{-a_2t}/a_2}{p_1/a_1+p_2/a_2}.\end{aligned} \tag{N11-13}$$

From the best fit of equation (N11-13) to the experimental data for the normalized fraction of immature proteins, the biexponential-distribution parameter values were determined as $a_1 = 10.0\,\text{h}^{-1}$, $a_2 = 2.31\,\text{h}^{-1}$, and $p_1 = 0.98$ for VenusNB (32 °C) and $a_1 = 6.26\,\text{h}^{-1}$, $a_2 = 2.60\,\text{h}^{-1}$, and $p_1 = 0.35$ for mCerulean (32 °C).

For the quantitative analysis of the IPTG-induced gene expression data (Fig. 3e), the gene activation occurs on a time scale far shorter than the super-Poisson, first gene expression burst process and is consistent with the fact that IPTG rapidly activates of the associated *tac* promoter by immediate inactivation of the *lac* repressor[25,26]. Accordingly, the distribution of the signal transduction time becomes equivalent to the distribution of the first protein burst time, i.e., $\hat{\varphi}_{st}(s) \cong \hat{\varphi}_b^{(1)}(s) = [1 + s\langle \tau_b^{(1)} \rangle \eta_{fb}^2]^{-1/\eta_{fb}^2}$ in the Laplace domain. Together with the burst-interval time distribution given by a gamma distribution, explicitly, $\hat{\varphi}_b(s) = [1 + s\langle \tau_b \rangle \eta_b^2]^{-1/\eta_b^2}$ in the Laplace domain, equations (N4-2) and (N4-9) read as

$$\langle \hat{R}(s) \rangle = \frac{\hat{\varphi}_{st}(s)}{1 - \hat{\varphi}_b(s)} \cong \frac{\hat{\varphi}_b^{(1)}(s)}{1 - \hat{\varphi}_b(s)} = \frac{[1 + s\langle \tau_b^{(1)} \rangle \eta_{fb}^2]^{-1/\eta_{fb}^2}}{1 - [1 + s\langle \tau_b \rangle \eta_b^2]^{-1/\eta_b^2}}. \quad \text{(N11-14a)}$$

$$\langle \hat{R}'(s) \rangle = \frac{\hat{\varphi}_b(s)}{1 - \hat{\varphi}_b(s)} = \frac{[1 + s\langle \tau_b \rangle \eta_b^2]^{-1/\eta_b^2}}{1 - [1 + s\langle \tau_b \rangle \eta_b^2]^{-1/\eta_b^2}}. \quad \text{(N11-14b)}$$

For the IPTG induction-response experiments at 37 °C, sfGFP was used as a fluorescent reporter. For numerical convenience, its maturation time distributions was assumed to be a hypo-exponential function, explicitly, equation (N11-2) with $n = 3$ and $C_i = a_i \prod_{j=1, j\neq i}^{3} a_j / (a_j - a_i)$, which well represents the time profile for the normalized fraction of immature proteins for sfGFP (Supplementary Fig. 2). For this maturation-time distribution model, the analytic expression of $S_m^{st}(t)$ is given by [see equation (N9-1) in Supplementary Note 9]

$$\begin{aligned} S_m^{st}(t) &= \frac{1}{\langle \tau_m \rangle}\int_t^\infty d\tau S_m(\tau) = \frac{1}{\langle \tau_m \rangle}\int_t^\infty d\tau \sum_{i=1}^{3} \frac{C_i}{a_i} e^{-a_i \tau} \\ &= \frac{\sum_{i=1}^{3} C_i e^{-a_i t} / a_i^2}{\sum_{i=1}^{3} C_i / a_i^2}. \end{aligned} \tag{N11-15}$$

From the best fit of equation (N11-15) to the experimental data for the normalized fraction of immature proteins, the hypo-exponential-distribution parameter values were determined as $a_1 = 6.92\,\mathrm{h}^{-1}$, $a_2 = 7.10\,\mathrm{h}^{-1}$, and $a_3 = 6.74\,\mathrm{h}^{-1}$ for sfGFP (37 °C).

Once models for $\langle \hat{R}(s) \rangle$, $\langle \hat{R}'(s) \rangle$, and $S_m^{st}(t)$ (or equivalently, $\varphi_m(t)$) are given, we can calculate the mean and variance of the mature protein number: equations (N11-1) and (N11-3) for $\langle \hat{n}(s) \rangle$ and equations (N11-5), (N11-6), (N11-8), and (N11-11) for $\mathcal{L}\left[\langle n^2(t) \rangle\right]$. For antibiotic stress-induced gene expression, $\langle \hat{R}(s) \rangle$, $\langle \hat{R}'(s) \rangle$, and $S_m^{st}(t)$ are given by equations (N11-12a), (N11-12b) and (N11-13), respectively. For IPTG-induced gene expression, $\langle \hat{R}(s) \rangle$, $\langle \hat{R}'(s) \rangle$, and $S_m^{st}(t)$ are given by equations (N11-14a), (N11-14b) and (N11-15), respectively. At time *t*, values of the mean and variance were then obtained through numerical Laplace inversion of $\langle \hat{n}(s) \rangle$ and $\mathcal{L}\left[\sigma_n^2(t)\right]$ using the Stehfest method[27].

For the quantitative analysis of the antibiotic stress-induced gene expression data (Fig. 3b, c), the theoretical results for the mean and variance of the mature protein number were multiplied by $c$ and $c^2$, respectively, where $c$ denotes the number-to-intensity conversion factor, based on the assumption of a linear relationship between the number of mature fluorescent proteins $n(t)$ and the corresponding fluorescence intensity $I(t)$, i.e.,

$I(t) = cn(t)$. For the quantitative analysis of the IPTG-induced gene expression data (Fig. 3e), the value of the protein degradation rate was fixed to $2.39\times10^{-1}$ $h^{-1}$, which was obtained from independent experiments for cell growth and intrinsic sfGFP degradation (Supplementary Fig. 4). Simultaneous optimization of the mean and variance was performed using the built-in MATLAB function *fmincon*. For both analyses, optimized parameter values are presented in Supplementary Tables 3 and 4.

### SUPPLYMENTARY FIGURES

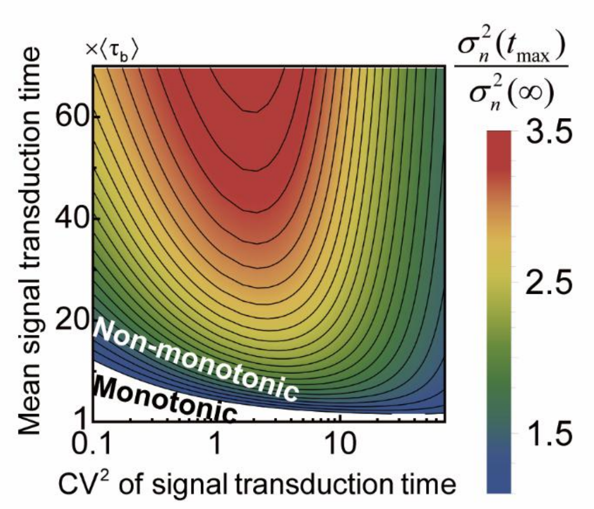


**Supplementary Figure 1. Dependence of the variance non-monotonicity parameter on mean and relative variance of signal transduction time.** The steady-state variance of mature protein levels as a function of the mean and relative variance (or $CV^2$) of the signal transduction time is calculated using the signal-induced adaptive gene expression model (Fig. 1a), with all model specifications identical to those used in Fig. 1 unless otherwise noted. The non-monotonic time dependence of the mature protein variance, $\sigma_n^2(t)$, is quantified by the ratio, $\sigma_n^2(t_{\max})/\sigma_n^2(\infty)$, between the maximum variance, $\sigma_n^2(t_{\max})$, attained over time and the steady-state variance, $\sigma_n^2(\infty)$. Ratio values greater than unity (shown in the heat map) indicate non-monotonic time dependence of the variance, whereas values smaller than unity (white background) indicate monotonic behavior. As shown in this figure, Fig. 1f corresponds to a crossover from monotonic to non-monotonic profiles as the relative variance of the signal transduction time increases from 0.1 to 1 and 10, while the mean signal transduction time is held fixed at $5\langle\tau_b\rangle$.

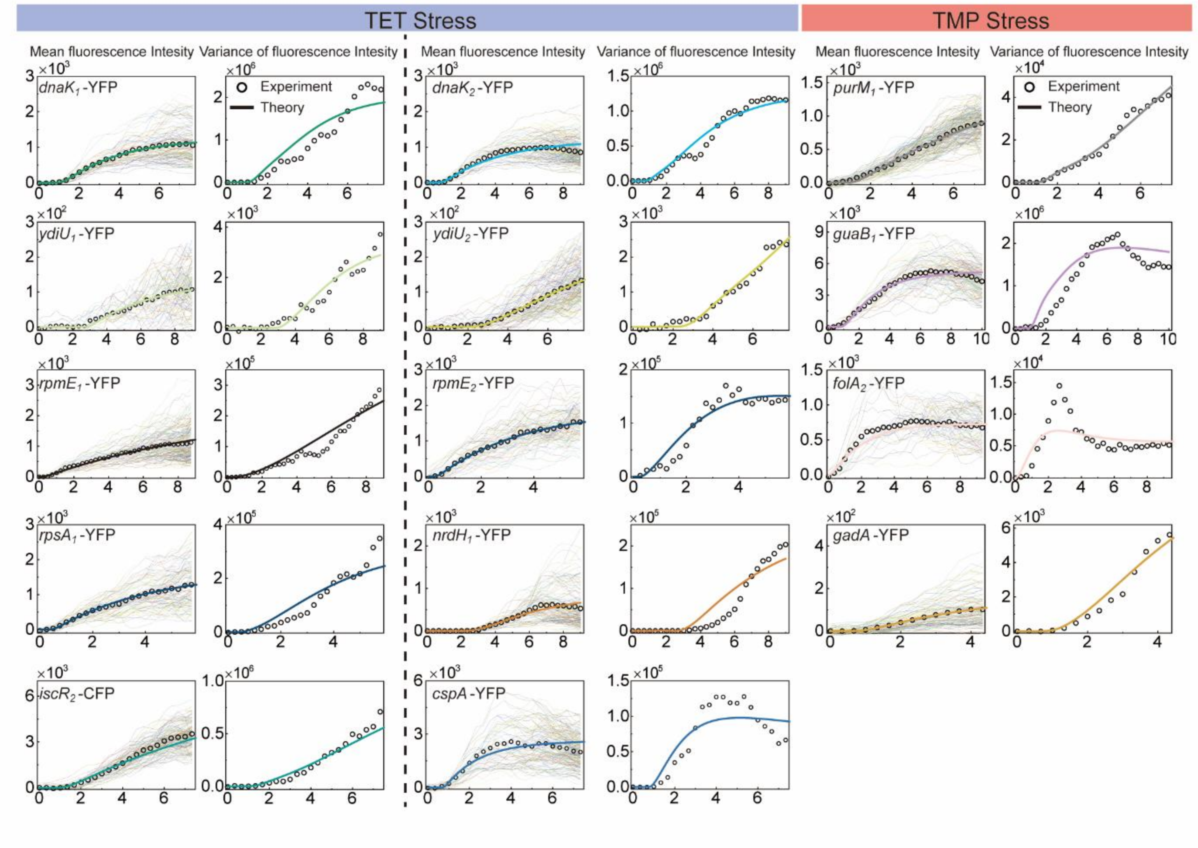


**Supplementary Figure 2. Quantitative analysis of antibiotic-induced bacterial gene expression.** Shown are the time-dependent mean and variance of reporter gene (*yfp* or *cfp*) expression levels in response to antibiotic stress by TET (left) and TMP (right). Subscript labels attached to promoter names indicate one of two independent datasets collected from different microcolonies. The symbols represent experimental results from single-cell gene expression time traces[22] (not shown in Fig. 3b, c of the main text) and the solid lines indicate the corresponding best fits of equations (1a) and (1b), each modified by a number-to-fluorescence intensity conversion factor (see Supplementary Note 10 and 11 for details of the quantitative analysis).

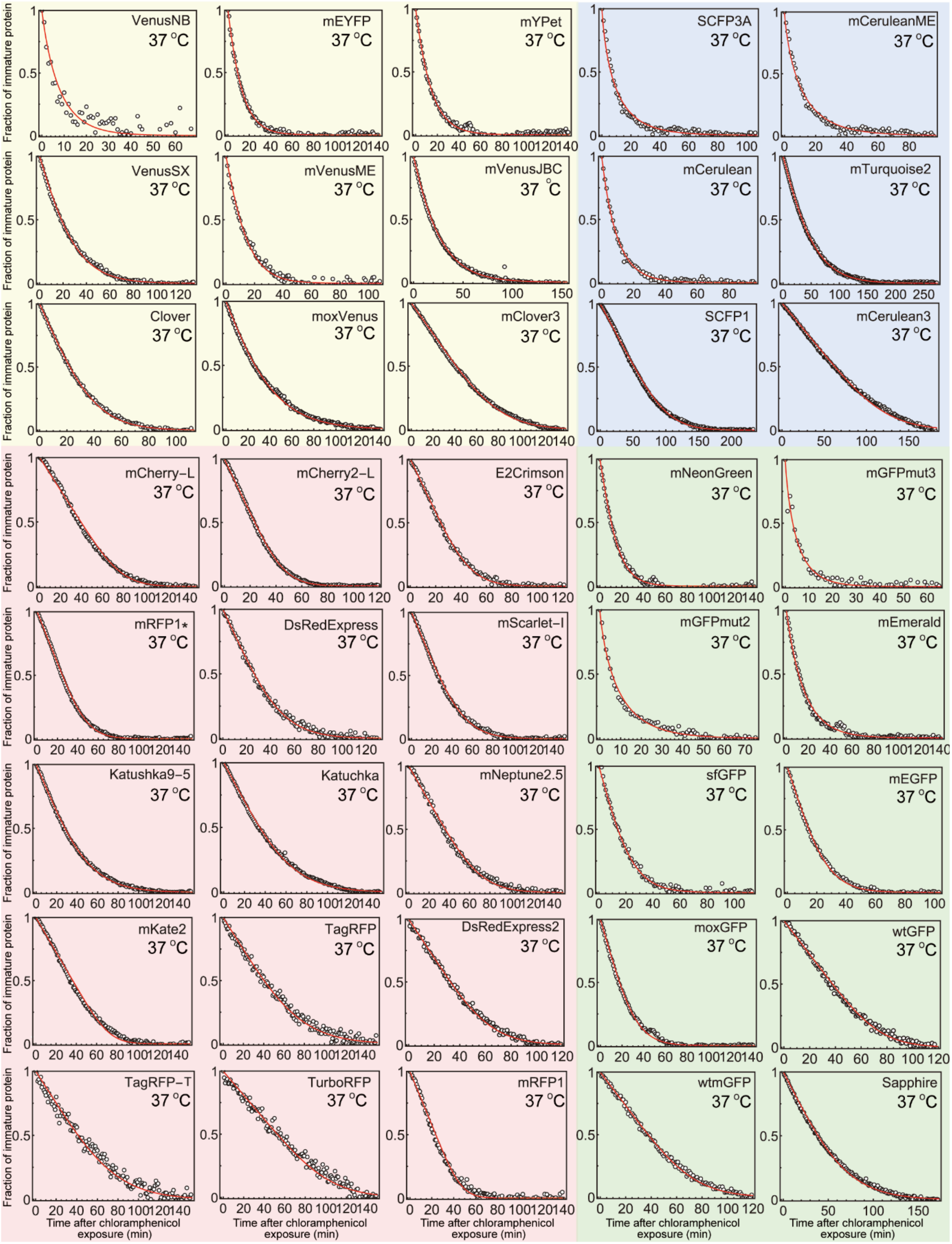
Fraction of immature protein
VenusNB
37 °C
mEYFP
37 °C
mYPet
37 °C
SCFP3A
37 °C
mCeruleanME
37 °C
VenusSX
37 °C
mVenusME
37 °C
mVenusJBC
37 °C
mCerulean
37 °C
mTurquoise2
37 °C
Clover
37 °C
moxVenus
37 °C
mClover3
37 °C
SCFP1
37 °C
mCerulean3
37 °C
mCherry-L
37 °C
mCherry2-L
37 °C
E2Crimson
37 °C
mNeonGreen
37 °C
mGFPmut3
37 °C
mRFP1*
37 °C
DsRedExpress
37 °C
mScarlet-I
37 °C
mGFPmut2
37 °C
mEmerald
37 °C
Katushka9-5
37 °C
Katuchka
37 °C
mNeptune2.5
37 °C
sfGFP
37 °C
mEGFP
37 °C
mKate2
37 °C
TagRFP
37 °C
DsRedExpress2
37 °C
moxGFP
37 °C
wtGFP
37 °C
TagRFP-T
37 °C
TurboRFP
37 °C
mRFP1
37 °C
wtmGFP
37 °C
Sapphire
37 °C
Time after chloramphenicol exposure (min)

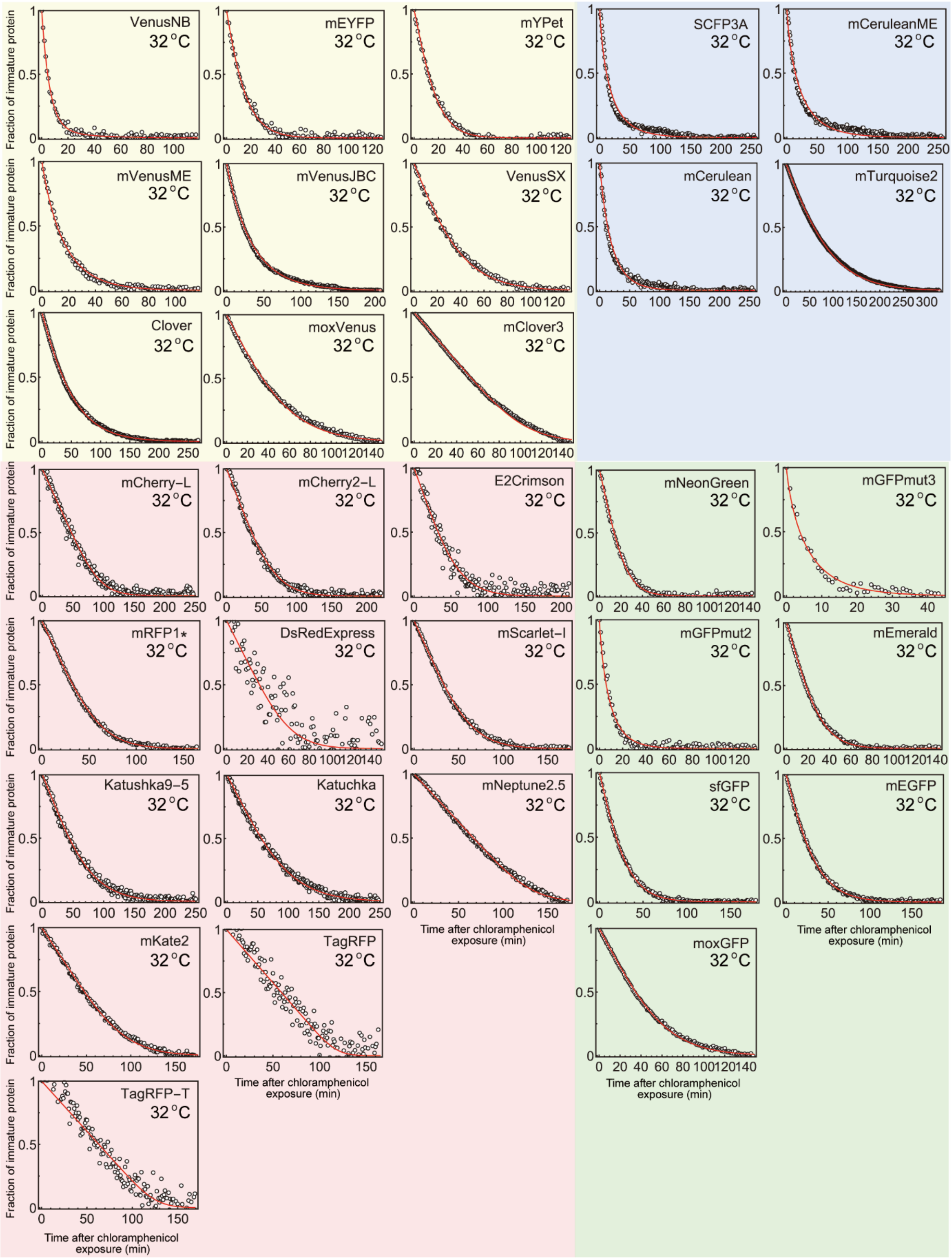


**Supplementary Figure 3. Cumulative distributions of maturation times for various fluorescent reporter proteins extracted from experimental data, along with their**

**corresponding model fits.** The cumulative maturation-time distribution for a given fluorescent protein was obtained from translation-stopped experiments[6]. Following chloramphenicol treatment of bacterial cells during gene expression, any subsequent increase in single-cell fluorescence intensity can be attributed to the maturation of proteins that were already synthesized prior to the treatment. These fluorescence intensity profiles were converted into the normalized fraction of immature proteins, shown here for 37 °C (upper figure) and 32 °C (lower figure). The shaded background in each panel indicates the fluorescence color of the corresponding fluorescent protein. The solid lines represent optimized model fits, assuming that maturation times follow a gamma distribution (see Supplementary Note 9 for details).

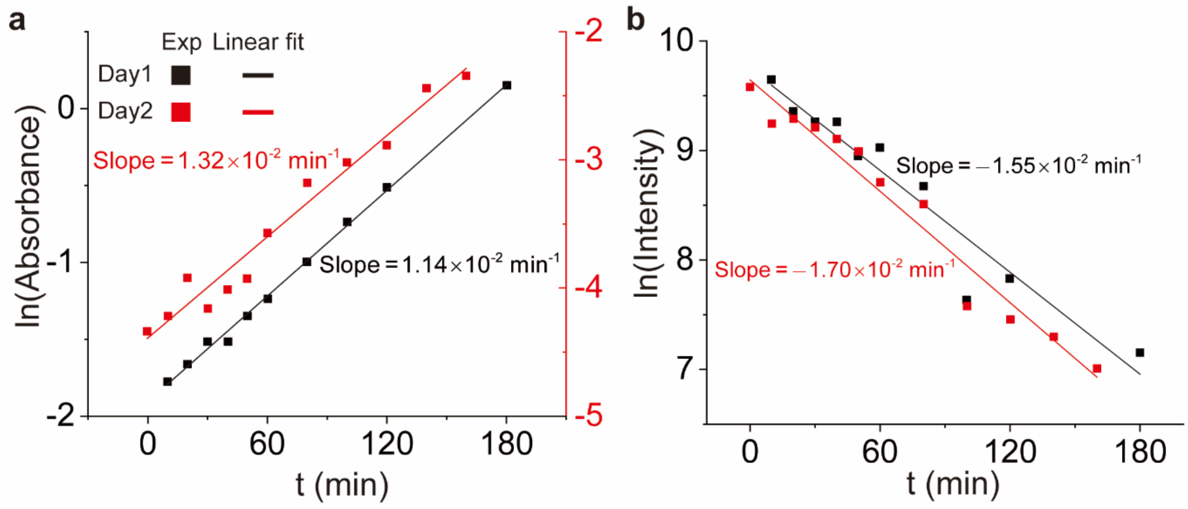


**Supplementary Figure 4. Experimental determination of the sfGFP degradation rate. a, b,** Cell growth and intrinsic sfGFP degradation profiles; optical density at 600 nm ($OD_{600}$) (a) and fluorescence intensity (b) were measured over time. The concentration of bacterial cells, estimated from $OD_{600}$ up to a proportionality constant, increases exponentially with time at a growth rate of $1.14\times10^{-2}$ min$^{-1}$ (Day 1) and $1.32\times10^{-2}$ min$^{-1}$ (Day 2). Under the non-inducing condition following IPTG removal (see Experimental Methods), the initial total amount of intracellular sfGFPs over bacterial cells, estimated from fluorescence intensity up to a proportionality constant, decreases exponentially with time at a decay rate of $1.55\times10^{-2}$ min$^{-1}$ (Day 1) and $1.70\times10^{-2}$ min$^{-1}$ (Day 2). The circles and lines represent experimental data and best exponential fits, respectively. Accounting for both dilution due to cell division and intrinsic protein degradation, we could determine the sfGFP degradation rate as the sum of the growth rate and the decay rate, each averaged over the two days. The value of the degradation rate is given by $2.39\times10^{-1}$ h$^{-1}$, which corresponds to the reciprocal of the mean protein lifetime, $1/\langle\tau_d\rangle$, when it is assumed that the distribution of protein lifetimes is given by an exponential distribution.

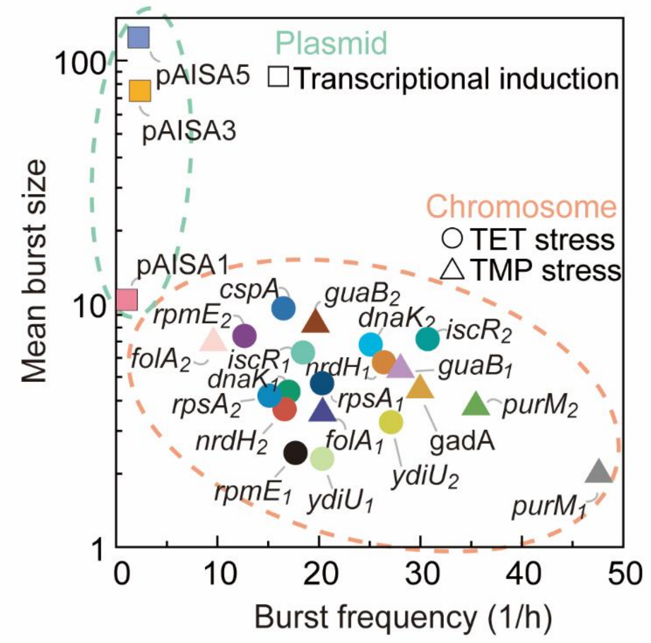


**Supplementary Figure 5. Effects of genetic platforms on protein bursts.** Shown are the mean burst size and burst frequency obtained from quantitative analyses of two experimental datasets (Supplementary Tables 3 and 4): antibiotic stress-induced expression of chromosome-integrated genes (circles: TET; triangles: TMP)[22] and IPTG-induced expression of plasmid-integrated genes (squares: UTR variants). Compared with plasmid-integrated genes (grouped by the green dashed line), chromosome-integrated genes exhibit higher burst frequencies and smaller burst sizes (grouped by the orange dashed line).

### SUPPLEMENTARY TABLES

**Supplementary Table 1.** Bacterial strains and plasmids used in this study

| | Description | Source |
|---|---|---|
| **Strains** | | |
| Mach1_T1R | $F^-$ φ80(lacZ)△M15 △lacX74 hsdR ($rK^-$ $mK^+$) △recA1398 endA1 tonA | Invitrogen |
| *E. coli* W3110 | *E. coli* $F^-$ $\lambda^-$ $rph^{-1}$ IN(rrnD, rrnE)1 | ATCC 27325 |
| EISA1 | *E. coli* W3110/pAISA1 | This study |
| EISA3 | *E. coli* W3110/pAISA3 | This study |
| EISA5 | *E. coli* W3110/pAISA5 | This study |
| **Plasmids** | | |
| pACYCDuet-1 | Expression vector, $Cm^R$, p15A ori | Novagen |
| pAISA0 | pACYC-$P_{Tac}$-*sfgfp*_$T_{BBa_B1006}$ | This study |
| pAISA1 | pACYC-$P_{Tac}$_$SynUTR_{sfgfp1}$-*sfgfp*_$T_{BBa_B1006}$ | This study |
| pAISA3 | pACYC-$P_{Tac}$_$SynUTR_{sfgfp3}$-*sfgfp*_$T_{BBa_B1006}$ | This study |
| pAISA5 | pACYC-$P_{Tac}$_$SynUTR_{sfgfp5}$-*sfgfp*_$T_{BBa_B1006}$ | This study |
| pQE80L-mK | pQE-80L-$P_{T5}$-*mkgsadh* | ref[28] |
| pQE80L-sfgfp | pQE-80L-$P_{T5}$-His6-*sfgfp* | This study |

**Supplementary Table 2.** Oligonucleotides used in this study.

| Name | Sequence (5′-3′) |
|---|---|
| pACYC_F | GCGGCCGCATTTCCTAATGCAG |
| pACYC_R | GCATGCCTGAAACCTCAGGCATTTG |
| Tac_R | GACGTCAATTGTTATCCGCTCACA |
| UTR1-Tac_F | TGTGAGCGGATAACAATTGACGTCACCGTAGGATTGATCGACAAT GAGCAAGGGCGAGGAG |
| UTR3-Tac_F | TGTGAGCGGATAACAATTGACGTCACAGGAGGATTGATCGTAGA TGAGCAAGGGCGAGGAG |
| UTR5-Tac_F | TGTGAGCGGATAACAATTGACGTCACAGGAGGATTGATCGAAAA TGAGCAAGGGCGAGGAG |
| sfgfp_F | CACCATCACCATCACGGATCCAGCAAGGGCGAGGAGCTG |
| sfgfp_R | GCATGGACGAGCTGTACAAGTGACTGCAGCCAAGCTTAATTAGCTG |
| pQE_F | GCATGGACGAGCTGTACAAGTGACTGCAGCCAAGCTTAATTAGCTG |
| pQE_R | CACCATCACCATCACGGATCCAGCAAGGGCGAGGAGCTG |

**Supplementary Table 3.** Optimized parameter values extracted from our quantitative analysis of the experimental data for antibiotic stress-induced gene expression (Figs. 3b, c, and Supplementary Fig. 3)[22].

| Promoter-reporter (upper section: TET stress; lower section: TMP stress) | Signal transduction | | | | Successive protein bursts | | | Protein decay rate | $c$ | $R^2$ |
|---|---|---|---|---|---|---|---|---|---|---|
| | Gene activation | | First protein burst | | Burst frequency | Burst size statistics | | | | |
| | $\langle t_{ga} \rangle$ (h) | $\eta^2_{ga}$ | $\langle \tau_b^{(1)} \rangle$ (h) | $\eta^2_{fb}$ | $1/\langle \tau_b \rangle$ ($h^{-1}$) | $\langle b \rangle$ | $\langle \delta b^2 \rangle$ | $1/\langle \tau_d \rangle$ ($h^{-1}$) | | |
| *dnaK₁*-YFP | 1.17 | 0.052 | 58.0 | 47.9 | 0.059 | 4.35 | 4.58 | 0.381 | 7.20 | 0.946 |
| *dnaK₂*-YFP | 0.858 | 0.050 | 57.6 | 70.1 | 0.040 | 6.79 | 6.86 | 0.300 | 2.30 | 0.957 |
| *cspA*-YFP | 0.673 | 0.065 | 4.87 | 21.4 | 0.061 | 9.57 | 10.3 | 0.698 | 12.7 | 0.860 |
| *ydiU₁*-YFP | 2.94 | 0.050 | 48.2 | 24.0 | 0.049 | 2.30 | 2.31 | 0.300 | 1.02 | 0.953 |
| *ydiU₂*-YFP | 2.86 | 0.050 | 18.4 | 44.7 | 0.037 | 3.26 | 3.28 | 0.100 | 0.445 | 0.986 |
| *nrdH₁*-YFP | 3.08 | 0.050 | 99.6 | 16.9 | 0.038 | 5.75 | 5.77 | 0.300 | 2.23 | 0.928 |
| *nrdH₂*-YFP | 3.08 | 0.050 | 1921 | 13.7 | 0.060 | 3.69 | 3.71 | 0.300 | 4.18 | 0.988 |
| *iscR₁*-CFP | 1.31 | 0.050 | 14.4 | 53.5 | 0.054 | 6.30 | 6.58 | 0.300 | 11.7 | 0.992 |
| *iscR₂*-CFP | 1.09 | 0.050 | 1.87 | 64.6 | 0.033 | 7.16 | 7.43 | 0.100 | 3.40 | 0.984 |
| *rpsA₁*-YFP | 0.486 | 0.051 | 23.2 | 35.1 | 0.049 | 4.71 | 4.73 | 0.300 | 5.74 | 0.951 |
| *rpsA₂*-YFP | 0.819 | 0.300 | 2.41 | 21.5 | 0.066 | 4.19 | 4.78 | 0.300 | 5.72 | 0.984 |
| *rpmE₁*-YFP | 0.246 | 0.192 | 69.0 | 36.6 | 0.057 | 2.44 | 2.44 | 0.138 | 6.28 | 0.953 |
| *rpmE₂*-YFP | 0.313 | 0.091 | 0.629 | 24.3 | 0.079 | 7.38 | 7.65 | 0.347 | 7.00 | 0.960 |
| *gadA*-YFP | 0.947 | 0.074 | 59.1 | 14.7 | 0.033 | 4.42 | 4.60 | 0.349 | 0.607 | 0.983 |
| *purM₁*-YFP | 1.25 | 0.050 | 16.5 | 124 | 0.021 | 1.99 | 1.99 | 0.100 | 2.22 | 0.989 |
| *purM₂*-YFP | 0.768 | 0.052 | 2.54 | 57.8 | 0.028 | 3.76 | 3.80 | 0.163 | 1.95 | 0.956 |
| *folA₁*-YFP | 0.500 | 0.077 | 90.6 | 59.5 | 0.049 | 3.55 | 3.59 | 0.622 | 6.51 | 0.921 |
| *folA₂*-YFP | 0.028 | 0.098 | 0.636 | 2.91 | 0.104 | 6.92 | 94.0 | 0.700 | 7.68 | 0.770 |
| *guaB₁*-YFP | 1.06 | 0.051 | 2.63 | 40.9 | 0.036 | 5.36 | 5.75 | 0.509 | 18.7 | 0.923 |
| *guaB₂*-YFP | 0.449 | 0.161 | 3.24 | 44.9 | 0.051 | 8.23 | 8.24 | 0.678 | 16.7 | 0.872 |

**Supplementary Table 4.** Optimized parameter values extracted from our quantitative analysis of the experimental data for transcriptional induction-induced gene expression (Fig. 3e).

| Plasmid | Signal transduction<br>First protein burst | | Successive protein bursts<br>Burst interval statistics | | Burst size statistics | | $R^2$ |
|---|---|---|---|---|---|---|---|
| | $\langle \tau_b^{(1)} \rangle$ (h) | $\eta_{fb}^2$ | $\langle \tau_b \rangle$ (h) | $\eta_b^2$ | $\langle b \rangle$ | $\langle \delta b^2 \rangle$ | |
| pAISA1 | 1.11 | 27.8 | 0.933 | 0.621 | 10.4 | 16.0 | 0.839 |
| pAISA3 | 43.1 | 23.0 | 0.426 | 0.015 | 75.1 | 227 | 0.884 |
| pAISA5 | 99.3 | 30.9 | 0.451 | 0.021 | 125 | 820 | 0.946 |
